# Conifers exhibit a characteristic inactivation of auxin to maintain tissue homeostasis

**DOI:** 10.1101/789420

**Authors:** Federica Brunoni, Silvio Collani, Rubén Casanova-Saéz, Jan Šimura, Michal Karady, Markus Schmid, Karin Ljung, Catherine Bellini

**Affiliations:** Umeå Plant Science Centre, Dept of Plant Physiology, Umeå University (Umu), Umeå, Sweden; Umeå Plant Science Centre, Dept of Forest Genetics and Plant Physiology, Swedish University of Agricultural Sciences (SLU), Umeå, Sweden; Institut Jean-Pierre Bourgin, UMR1318 INRA-AgroParisTech Versailles, France

**Keywords:** auxin conjugates, auxin homeostasis, conifers, *GH3* genes, indol-3-acetic acid (IAA), *Picea abies*

## Abstract

- Dynamic regulation of the levels of the natural auxin, indol-3-acetic acid (IAA), is essential to coordinate most of the physiological and developmental processes and responses to environmental changes. Oxidation of IAA is a major pathway to control auxin concentrations in Arabidopsis and, along with IAA conjugation, to respond to perturbation of IAA homeostasis. However, these regulatory mechanisms are still poorly investigated in conifers. To reduce this gap of knowledge, we investigated the different contribution of the IAA inactivation pathways in conifers.
- Mass spectrometry-based quantification of IAA metabolites under steady state conditions and after perturbation was investigated to evaluate IAA homeostasis in conifers. Putative *Picea abies GH3* genes (*PaGH3*) were identified by a comprehensive phylogenetic analysis including Arabidopsis and basal land plants. Auxin-inducible *PaGH3* genes were identified by expression analysis and their IAA-conjugating activity was explored.
- Compared to Arabidopsis, oxidative and conjugative pathways differentially contribute to reduce IAA levels in conifers. We demonstrated that the oxidation pathway plays a marginal role in controlling IAA homeostasis in spruce. On the other hand, an excess of IAA rapidly activates GH3-mediated irreversible conjugation pathways.
- Taken together, these data indicate that a diversification of IAA inactivation mechanisms evolved specifically in conifers.

## Introduction

The phytohormone auxin mediates a variety of different developmental processes during a plant’s life cycle, acting also as an intercellular signal integrating environmental inputs and growth responses. An important level of regulation in auxin responses is the establishment of concentration gradients between cells and tissues (Vanneste & Friml, 2009; Ljung, 2013). Thus, it is critical for plants to tightly control differential auxin distribution, both at spatial and temporal levels. Metabolic and transport mechanisms jointly generate auxin gradients within tissues (Ruiz Rosquete *et al.*, 2012; Ljung, 2013). Auxin transport depends on the differential localization of influx and efflux carriers at the plasma membrane, providing the plant with a carrier-driven mechanism that controls auxin directionality and distribution within tissues (Vanneste & Friml, 2009). Additionally, auxin levels are regulated via the balance between the rates of auxin biosynthesis and auxin inactivation, extending the complexity of the regulatory network of cellular auxin homeostasis (Ljung, 2013; Kramer & Ackelsberg, 2015). The predominant auxin found in plants is indole-3-acetic acid (IAA). Multiple auxin biosynthetic pathways that rely on tryptophan (Trp) as a precursor have been proposed. The elucidation of these biosynthetic pathways demonstrated that IAA is locally produced and not ubiquitously, as it was earlier assumed (Normanly, 2010; LeClere *et al.*, 2002; Zhao, 2018). Evidences for a Trp-independent pathway for IAA biosynthesis also exist but this pathway is less understood (Ljung, 2013; Zhao, 2018). The reduction of the cellular levels of free IAA also relies on auxin inactivation mechanisms, such as conjugation and degradation. IAA can be conjugated to various amino acid, peptide, and sugar moieties. Auxin conjugates might function as short-term intermediates that can release free IAA upon hydrolysis when required (LeClere *et al.*, 2002; Ludwig-Müller, 2011). IAA can be reversibly conjugated *via* ester linkages to glucose by UDP-glucosyl transferases to produce indole-3-acetyl-1-glucosyl ester (IAGlc) (Jackson *et al.*, 2001). Members of the GRETCHEN HAGEN3 (GH3) family of acyl amido synthetases mediate conjugation of IAA with amino acids (Staswick *et al.*, 2005; Westfall *et al.*, 2011). IAA amino acid conjugates are regarded as either reversible or irreversible IAA metabolites based on *in vitro* activity and *in planta* feeding assays (Östin *et al.*, 1998; Kowalczyk & Sandberg, 2001). Conjugation to particular moieties, such as aspartate (Asp) and glutamate (Glu) to produce indole-3-acetyl-L-aspartic acid (IAAsp) and indole-3-acetyl glutamic acid (IAGlu), seems to lead to degradation (Östin *et al.*, 1998; Tam *et al.*, 2000), whereas IAA conjugates with other amino acids (*e. g.*, alanine, leucine or phenylalanine) are transient storage compounds that could be hydrolysed back to free IAA *via* auxin amino acid conjugate hydrolases (Kowalczyk & Sandberg, 2001; LeClere *et al.*, 2002). The oxidation of IAA into oxindole-3-acetic acid (oxIAA) is one of the major catabolic pathways to inactivate auxin (Östin *et al.*, 1998; Peer *et al.*, 2013; Pěnčík *et al.*, 2013). Recent findings demonstrated that the DIOXYGENASE FOR AUXIN OXIDATION (DAO) protein, a 2-oxoglutarate-dependent-Fe (II) dioxygenase, catalyzes conversion of IAA to oxIAA (Zhao *et al.*, 2013; Porco *et al.*, 2016; Zhang *et al.*, 2016). oxIAA can be further glucosylated to oxindole-3-acetyl-1-glucosyl ester (oxIAGlc) (Östin *et al.*, 1998; Kai *et al.*, 2007). The characterization of these metabolic pathways together with the identification of the key components of the auxin conjugation and degradation machineries have significantly improved our understanding of their respective contribution to the regulation of auxin homeostasis. For example, metabolic profiling of the loss-of-function *dao1* mutant showed that, despite the reduction of oxIAA concentration in the mutant, no significant accumulation of IAA was observed (Porco *et al.*, 2016). Very interestingly, the levels of IAA conjugates, such as IAAsp and IAGlu, were found to be much higher in the *dao1* mutant than in wild-type plants and the accumulation of these IAA metabolites was associated with an increase of *GH3* gene expression in the *dao1* mutant (Porco *et al.*, 2016). These findings demonstrated that *DAO1* acts in concert with *GH3* genes to maintain optimal auxin levels, and thus the regulation of IAA homeostasis by these auxin inactivation pathways is redundant (Stepanova & Alonso, 2016; Zhang & Peer, 2017). Additionally, differences in enzyme kinetics and expression levels between DAO and GH3 suggested that DAO, having much slower enzyme kinetics compared with GH3 proteins, contributes to maintain constitutively basal auxin levels under normal growth conditions, while GH3 rapidly responds to environmental factors that increase cellular IAA levels (Mellor *et al.*, 2016; Stepanova & Alonso, 2016). Although the majority of the knowledge about auxin metabolism is based on *Arabidopsis thaliana* studies, the conjugation and oxidation mechanisms seem to be present also in many other species (Cooke *et al.*, 2002; Ludwig-Müller, 2011; Zhang & Peer, 2017). Analysis of endogenous IAA metabolites in cyanobacteria, algae and bryophytes revealed that free IAA and its primary catabolite oxIAA contribute the most to the total auxin pool in these species (Drábková *et al.*, 2015; Žižková *et al.*, 2017). IAAsp concentration was close to the detection limit in algae and cyanobacteria under normal growth conditions, while exogenous application of radiolabeled IAA led to accumulation of IAAsp and IAGlc in green algae (Žižková *et al.*, 2017). IAA amino acid conjugates were not found abundant in bryophytes and only IAAsp and IAGlu were detected in some species, if present at all (Drábková *et al.*, 2015). Ester-linked conjugates, such as IAGlc and oxIAGlc, were also present in liverworts and mosses but they did not contribute considerably to the total pool of auxin (Drábková *et al.*, 2015). In gymnosperms, it has been postulated that the two different categories of IAA conjugates have a different function during early growth of Scots pine (*Pinus sylvestris*) seedlings (Ljung *et al.*, 2001). Ester-linked conjugates of IAA were hydrolyzed during seed germination acting as a source for release of free IAA, while IAAsp, together with oxIAA, contributed the less to the IAA pool size in the seed but progressively accumulated during initial seedling growth (Ljung *et al.*, 2001). These findings suggest that flowering plants did not evolve *de novo* mechanisms for the homeostatic control of the IAA pool size, but have rather developed mechanisms from metabolic pathways that were already present in the early land plants and algae.

However, the function of these processes and the study of specific enzymes catalyzing these reactions in non-model species remain poorly understood. Until recently, this information was inaccessible due to unavailable genomic resources and the lack of suitable tools for the quantification of auxin metabolites. The release of conifers’ draft genomes (Birol *et al.*, 2013; Nystedt *et al.*, 2013; Neale *et al.*, 2014) combined with the development of sensitive methods for the identification and quantification of auxin metabolites, have opened new possibilities for functional studies about the IAA homeostatic mechanisms in conifers. This article provides strong evidence that the formation of oxIAA does not contribute considerably to maintain IAA homestasis in Norway spruce (*Picea abies*), as well as in other conifers such as Scots pine and Logdepole pine (*Pinus contorta*) in contrast to Arabidopsis. On the other hand, the irreversible conversion of IAA to IAAsp and IAGlu seems to act constitutively in steady state conditions and to be the main pathway induced in response to perturbation of the IAA content, to maintain IAA homeostasis in conifers. We also report the identification of auxin-inducible Norway spruce Group II members of the *GH3* family and confirm that the corresponding recombinant proteins catalyze the conjugation of IAA with Asp and/or Glu. Taken together, our data suggest that the sophisticated regulatory machinery controlling IAA homeostasis is conserved in conifers although the respective contribution of the oxidative and conjugation pathways is different compared to angiosperms. This suggests a diversification of the homeostatic control of IAA in gymnosperms.

## Materials and methods

### Plant material and growth conditions

Norway spruce seeds (*Picea abies* L. Karst) were provided by Sveaskog (Lagan, Sweden) and pine seeds (*Pinus sylvestris* L. and *Pinus contorta* Dougl.) were provided by SkogForsk (Sävar, Sweden). Seeds were soaked in tap water for 12 h at 4°C and then sown in fine wet vermiculite. Germination and seedlings growth occurred in a growth chamber at light intensity of 150 μE m^-2^ s^-1^ under long day condition (16 h light and 8 h dark). Temperatures were set to 22°C during the light and 18°C during the night.

### IAA metabolite profiling and feeding experiment with unlabeled and labeled IAA

Spruce and pine seedlings were sampled 14 days after sowing and dissected organs (cotyledons, hypocotyl and root) were collected in four independent replicates (10 mg of tissue per sample). For feeding experiments with unlabeled IAA, 2-week-old spruce and pine seedlings were first cultured in flasks containing 50 ml of half-strength (½) Murashige-Skoog (MS, Duchefa, Harleem, The Netherlands) liquid medium for 24 h under gentle shaking and in darkness. Liquid cultures were subsequently supplemented with 10 μM IAA for 0, 6 and 24 h under gentle shaking and in darkness. Mock treated seedlings were used as control. For each time point, 10 mg of roots were collected in four independent replicates. For feeding experiments with labeled IAA, 2-week-old spruce seedlings were first incubated in ½ MS liquid medium for 24 h under gentle shaking and in darkness. Liquid cultures were subsequently supplemented with 10 μM [^13^C_6_]-IAA for 0, 6 and 24 h under gentle shaking and in darkness. For each time point, 10 mg of spruce roots were collected in five independent replicates. The extraction, purification and LC-MS/MS analysis of endogenous concentrations of auxin and its metabolites were carried out according to Novák *et al.* (2012). Briefly, approx. 10 mg of frozen material per sample was homogenized using a bead mill (27 hz, 10 min, 4°C; MixerMill, Retsch GmbH, Haan, Germany) and extracted in 1 ml of 50 mM sodium phosphate buffer containing 1% sodium diethyldithiocarbamate and a mixture of ^13^C_6_- or deuterium-labeled internal standards. After centrifugation (14000 RPM, 15 min, 4°C), the supernatant was transferred into new Eppendorf tubes. Afterwards, the pH was adjusted to 2.5 by 1 M HCl and applied on preconditioned solid-phase extraction columns (Oasis HLB, 30 mg 1 cc, Waters Inc., Milford, MA, USA). After sample application, the column was rinsed with 2 ml 5% methanol. Compounds of interest were then eluted with 2 ml 80% methanol. Mass spectrometry analysis and quantification were performed using a LC-MS/MS system comprising of a 1290 Infinity Binary LC System coupled to a 6490 Triple Quad LC/MS System with Jet Stream and Dual Ion Funnel technologies (Agilent Technologies, Santa Clara, CA, USA).

### Sequences and phylogenetic analysis

The gene family information available in PlantGenIE (https://plantgenie.org; Sundell *et al.*, 2015) was used to retrieve putative *GH3* genes from *P. abies* genome. The same procedure was adopted to retrieve GH3-like protein sequences also from non-seed plants, such as *Selaginella moellendorffii* and *Physcomitrella patens. A. thaliana* GH3 proteins were used to blast embryophyte and chlorophyte genomes in Phytozome v12.1 (https://phytozome.jgi.doe.gov/pz/portal.html; Goodstein *et al.*, 2012), allowing the selection of additional putative GH3 protein sequences from *Marchantia polymorpha* and *Sphagnum fallax*, since no GH3 sequences were found in chlorophytes. All the putative GH3 proteins identified in this initial search were examined by manual curation of protein motif scan using Pfam (http://pfam.xfam.org/; El-Gebali *et al.*, 2019) for the GH3 motif (Pfam id: 03321; GH3 auxin-responsive promoter). A total of 14 *P. abies*, 21 *S. moellendorffii*, 2 *P. patens*, 3 *S. phallax* and 2 *M. polymorpha* were sorted as unique sequences that contained the conserved protein motif of interest. The available sequence information of all sorted and predicted proteins (both full-length and partial), the corrected spruce sequences information obtained in our laboratory (**Fig. S1**) and Arabidopsis GH3 protein sequences were used for the phylogenetic analysis. Multiple sequence alignment was performed using MUSCLE (Edgar, 2004) as implemented in MEGA6 (http://megasoftware.net/; Tamura *et al.*, 2013) under default parameters. The phylogenetic trees were constructed with the Neighbor-Joining method (Saitou & Nei, 1987) in MEGA6 with 1000 bootstrap iterations. The evolutionary distances were computed using the Dayhoff matrix based method (Schwarz & Dayhoff, 1979) and are reported in number of amino acid substitutions per site.

### RNA isolation and real-time PCR

Root samples for analyzing the mRNA levels of *PaGH3*s in response to auxin, were collected from spruce seedlings that were cultured in flasks containing 50 ml of ½ MS liquid media supplemented with 10 μM IAA for 0 or 6 hours under gentle shaking and in darkness. Mock treated seedlings were used as a control. Samples were frozen in liquid nitrogen and stored at −80°C after collection. Total RNA was isolated using the Spectrum Plant Total RNA kit (Sigma-Aldrich, USA) according to the manufacturer’s instructions. For each sample, 1 µg of total RNA was reverse transcribed with iScript cDNA Synthesis Kit (Bio-rad, USA) according to manufacturer’s instructions. Quantitative real-rime PCR (qRT-PCR) was performed on a CFX384 Touch Real-Time PCR Detection System (Bio-Rad Laboratories, USA) using 2x LightCycler 480 SYBR Green I Master (Roche, Germany). The three step cycling program was as follows: 95°C for 3 min, followed by 40 cycles at 95°C for 10s, 60°C for 15s and 72°C for 30s. The melting curve analysis was conducted between 65°C-95°C. The specificity of the PCR products was confirmed by analyzing melting curves and sequencing. Only primer pairs that produced a linear amplification and qPCR products with a single-peak melting curves were used for further analysis. The efficiency of each pair of primers was determined from the data of amplification Cycle threshold (Ct) value plot with a serial dilution of mixture cDNA and the equation E=10^(−1/slope)^-1. *P. abies eIF4A* gene (MA_50378g0010) was used as a constitutive internal standard, which showed no clear changes in Ct values, to normalize the obtained gene expression results. Expression levels were calculated using the ΔΔCt method (Pfaffl, 2001). Three biological replicates, each with three technical replicates were performed for each test. Primer sequences are listed in **Table S1**.

### 5’- and 3’-RACE, cloning, protein expression and enzyme assay

The full-length coding regions of *PaGH3.16, PaGH3.17, PaGH3.gII.8* and *PaGH3.gII.9* were obtained by combining incomplete gene annotation from Norway spruce genome (https://congenie.org; Nystedt *et al.*, 2013), with 5’- and 3’-RACE (Gene Racer Kit, Invitrogen, USA), which was carried out according to the manufacturer’s instructions. The full-length coding regions were then amplified by PCR from spruce cDNA template using gene-specific primers (**Fig. S1; Table S1**) with additional *XbaI* and *HindIII* or *NheI* sites required to clone into pETM11 expression vector. Plasmids expressing recombinant spruce proteins were transformed into *Escherichia coli* BL21 (DE3). Recombinant proteins production was induced by the addition of 0.1 mM isopropyl-D-thiogalactopyrano-side (IPTG). Cells were grown over night at 20°C with constant shaking at 250 rpm. Protein expression was tested by Western blot with anti-6x His antibody horseradish peroxidase (HPR) conjugate (Sigma A7058-1VL, 1:10000 dilution). IAA conjugation assays were performed as described in Brunoni *et al.* (2019).

## Results

### 1. IAA conjugation processes mostly control IAA homeostasis in conifer seedlings

To investigate how IAA is metabolized in conifers, basal IAA metabolism was analyzed in three different conifer species, *Picea abies, Pinus sylvestris* and *Pinus contorta*, under steady state growth conditions. Vermiculite-grown conifer seedlings were harvested 14 days after germination and profiling of endogenous IAA metabolites was carried out in cotyledons, hypocotyl and root, separately. **Fig. 1** and **Fig. S2** show IAA metabolites’ profiling in the different organs of spruce and pine seedlings, respectively. For all conifers, free IAA was abundant in roots while the concentration was lower in the cotyledons and hypocotyl and, together with IAGlc, contributed to most of the total auxin metabolite pool in both spruce and pine seedlings (**Fig. 1; Fig. S2**). Concerning the irreversible IAA catabolism, IAAsp and IAGlu were the most abundant IAA catabolites while oxIAA occurred at very low concentration in all spruce or pine organs analyzed (**Fig. 1; Fig. S2**). Interestingly IAAsp and IAGlu were not equally distributed throughout the pine organs, with IAAsp levels close to limit of detection in the hypocotyl of *P. sylvestris* and IAGlu not detected in roots or close to the limit of detection in the hypocotyl of *P. contorta* (**Fig. S2**). The glucosylester oxIAGlc is present at high levels in Arabidopsis seedlings (Kai *et al.*, 2007; Porco *et al.*, 2016) and was suggested to be synthesized *via* glucosylation of oxIAA and not oxidation of IAGlc (Tanaka *et al.*, 2014; Porco *et al.*, 2016). Compared to Arabidopsis seedlings, oxIAGlc was found to be much less abundant in all conifer seedlings (**Fig. 1; Fig. S2**). oxIAGlc was detected at low level in all three organs from pine seedlings, but it was below the detection limit in cotyledons from spruce seedlings (**Fig. 1; Fig. S2**). Taken together, these results suggest that, under steady state conditions, the conjugation pathways are likely to be the main contributors to IAA homeostasis compared to the formation of oxIAA and oxIAGlc in conifers.

**Figure 1.**
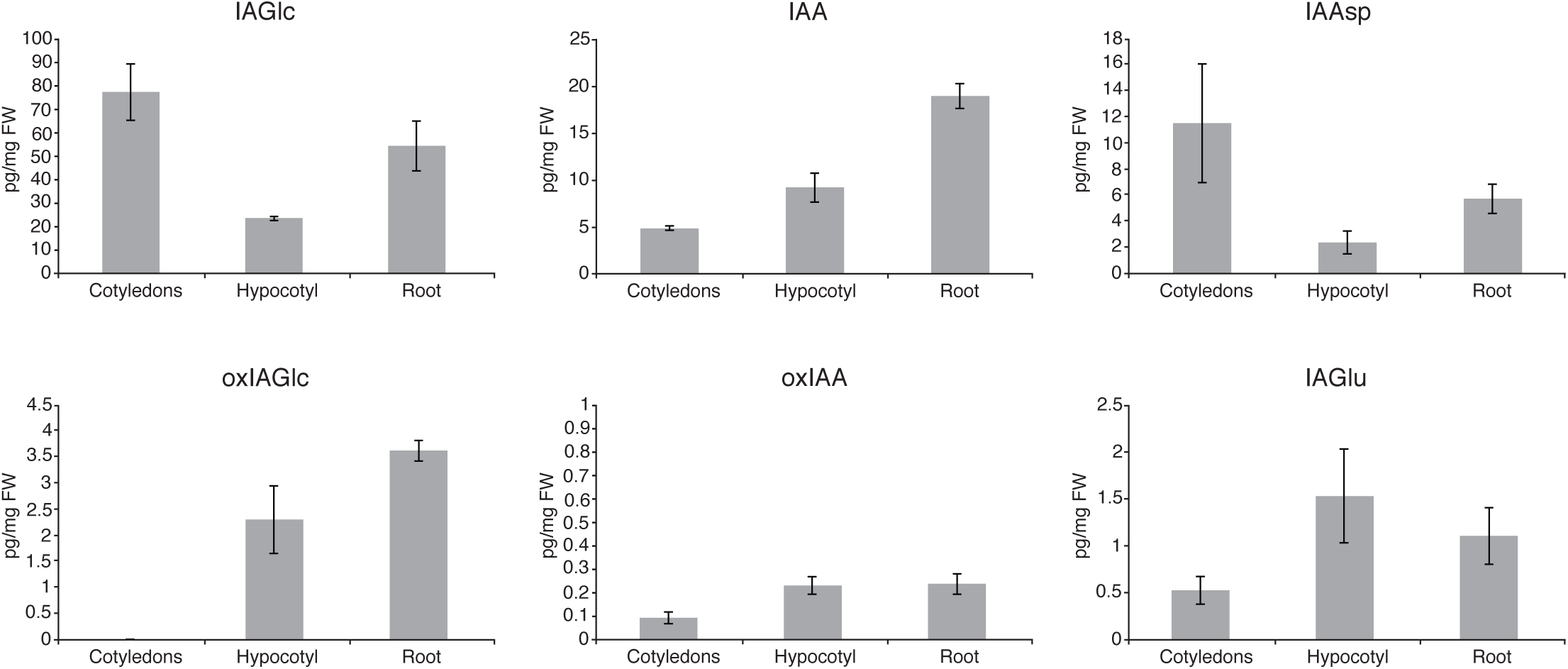
Levels of IAA metabolites in different organs of *Picea abies* seedlings. IAA and IAA metabolites oxIAA, oxIAGlc, IAGlc, IAAsp and IAGlu were quantified in cotyledons, hypocotyl and root from 2-week-old spruce seedlings. The level of oxIAGlc in cotyledons was under the detection limit of the used LC-MS/MS method. The concentrations of all metabolites are in picograms per milligram fresh weight (FW). Error bar indicates ±SD (*n* = 4).

### 2. Exogenous application of IAA rapidly activates conjugation pathway in conifer roots

Application of exogenous IAA and generation of transgenic or mutated lines, in which the functionality of key regulators in auxin metabolism are affected, lead to perturbations of IAA homeostasis (Staswick *et al.*, 2005; Park *et al.*, 2007; Zao *et al.*, 2013; Pěnčík *et al.*, 2013; Porco *et al.*, 2016). When mutant or transgenic lines are not easily available, as with spruce, metabolic analysis of IAA-treated plants represents a valid alternative to study the relative contribution of the different inactivation mechanisms to the regulation of IAA levels. When spruce seedlings were treated with 1 or 10 µM of exogenous IAA, alterations in the levels of IAA metabolites was similar in the 3 different organs (cotyledons, hypocotyl and root), but more evident in root tissues (**Fig. S3**), and 10 µM of IAA was the most effective concentration to be used (**Fig. S3**).

We then analyzed the effect of disturbed IAA homeostasis by incubating 2-week-old spruce and pine seedlings with 10 µM IAA, and roots were harvested at 6 and 24 hours after treatment. After 6 hours, exogenous auxin application led to a strong endogenous IAA accumulation but after 24 hours free IAA levels dropped dramatically, reaching the basal levels detected in mock treatments, suggesting that IAA inactivation machinery was activated within this time span (**Fig. 2a; Fig. S4**). Activation of the IAA metabolism occurred in all 3 species already after 6 hours, and all IAA metabolites increased dramatically after treatment with exogenous IAA compared to the levels detected in mock treatments (**Fig. 2a; Fig. S4**). Among the primary degradation products, IAAsp and IAGlu were much higher than oxIAA, and accumulated over time in spruce (**Fig. 2a**). IAA was more efficiently conjugated with Asp than with Glu, as much higher levels of IAAsp than IAGlu were measured (**Fig. 2a**). The levels of the ester-linked conjugates IAGlc was also higher in IAA treated seedlings than in mock treated ones at 6 and 24 hours, suggesting that glucosylation of IAA also contributed to the homeostatic regulation (**Fig. 2a**). After exogenous IAA application, the amount of oxIAGlc was also increasing, but the levels were lower more than 100-fold than IAGlc, confirming the marginal role of the oxidative pathways in homeostatic regulation of IAA in spruce (**Fig. 2a**). A similar metabolic pattern was observed in pine roots treated with exogenous auxin (**Fig. S4**). Taken together, these results indicate that the irreversible conjugation to form IAAsp and IAGlu was rapidly activated by increased free IAA levels in conifer seedlings.

**Figure 2.**
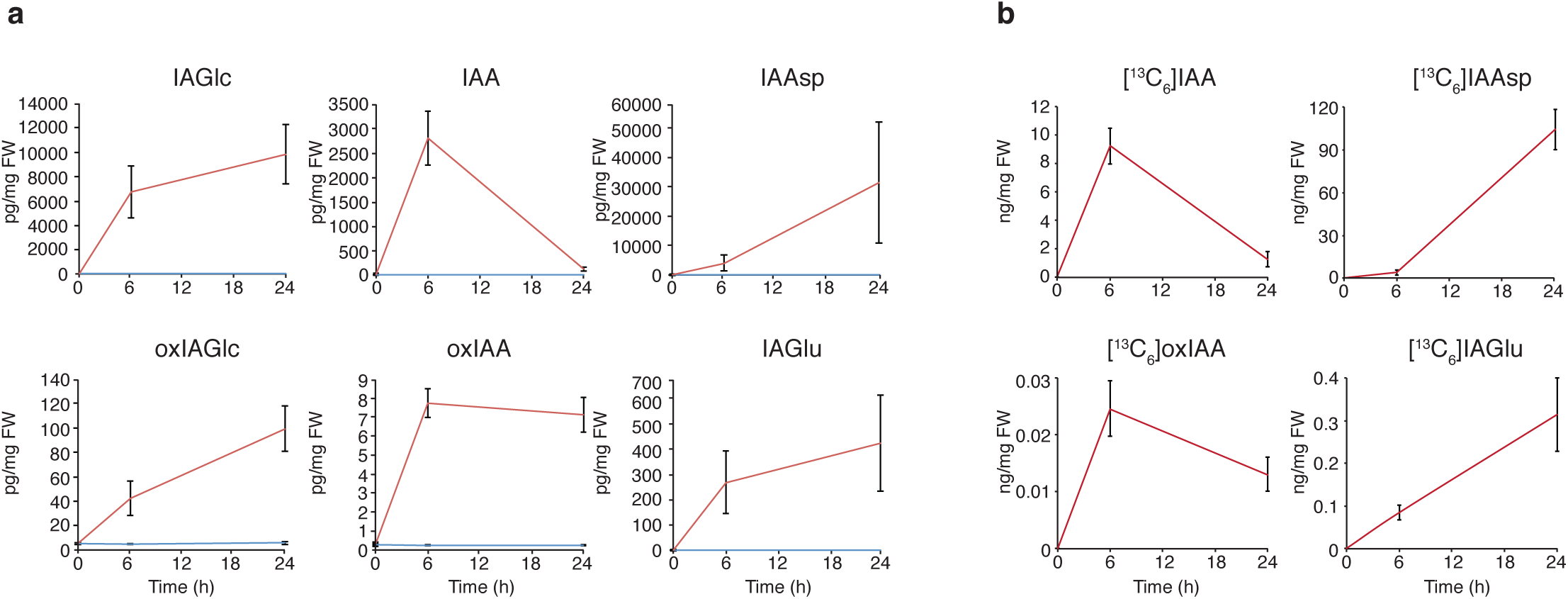
Concentrations of IAA metabolites in *Picea abies* roots after feeding with unlabeled or labeled IAA. (**a**) Two-week-old spruce seedlings were incubated with (red line) or without (blue line) 10 μM unlabeled IAA and the levels of IAA and IAA metabolites oxIAA, oxIAGlc, IAGlc, IAAsp and IAGlu were quantified in roots after different incubation times. The concentrations of all metabolites are in picograms per milligram fresh weight (FW). (**b**) Two-week-old spruce seedlings were incubated with 10 μM [^13^C_6_]IAA and the levels of [^13^C_6_]-oxIAA, [^13^C_6_]-IAAsp and [^13^C_6_]-IAGlu were quantified in roots after different incubation times. The concentrations of all metabolites are in nanograms per milligram fresh weight (FW). Error bars indicate ±SD (*n* = 4 or *n* = 5, respectively).

To study the rates of IAA degradation in spruce roots, we then performed a feeding experiment in which [^13^C_6_]-IAA was applied to whole seedlings and the degradation of IAA was measured by quantification of labeled IAA metabolites in roots. Feeding labeled IAA resulted in the formation of mainly [^13^C_6_]-IAAsp, low levels of [^13^C_6_]-IAGlu and almost no [^13^C_6_]-oxIAA after 6 or 24 hours (**Fig. 2b**), in accordance with the results obtained with unlabeled IAA (**Fig. 2a**). These findings indicate that the high endogenous levels of IAAsp and IAGlu observed in spruce originated from *de novo* synthesis.

### 3. The spruce genome has several *GH3* genes that belong to Group II of the *GH3* family

Our metabolic data have highlighted the importance of irreversible conjugative pathways for the regulation of IAA homeostasis in conifers. Auxin-conjugating enzymes of Group II of the GH3 family mediate the formation of IAAsp and IAGlu in Arabidopsis and this mechanism seems to be highly conserved all over the plant kingdom (Staswick *et al.*, 2005; Terol *et al*., 2006; Reddy *et al.*, 2006; Ludwig-Müller *et al.*, 2009; Ludwig-Müller, 2011). To identify the genes responsible for the conjugation of IAA with amino acids in conifers, we have analyzed the GH3 family of IAA-amido synthases and used *P. abies* as model species due to the availability of genomic resources (Nystedt *et al.*, 2013). Putative *GH3* genes of *P. abies* were retrieved using the gene family information present in PlantGenIE (https://plantgenie.org; Sundell *et al.*, 2015). This allows the identification of 14 *P. abies* genes (henceforth referred to as *PaGH3* genes) encoding full-length or partial GH3 proteins (**Fig. 3**). We named these genes according to their putative orthologs in Arabidopsis, as explained below. Only 5 *PaGH3* genes encode full-length proteins (PaGH3.17, PaGH3.16, PaGH3.gII.1, PaGH3.gII.8 and PaGH3.gII.9), while the remaining 9 are partial genes, lacking a portion in either the amino-, the carboxy- or both terminal regions. Three conserved sequence motifs involved in ATP/AMP binding that are characteristic of the acyl-adenylate/thioester-forming enzyme superfamily were previously identified and shown to be also conserved in the GH3 family (Staswick *et al.*, 2002; Terol *et al.*, 2006). The multiple alignment with the available deduced amino acid GH3 sequences from *P. abies* and Arabidopsis showed that all 5 full-length proteins, together with the partial protein sequence from PaGH3.gII.7, contain the 3 motifs, the partial proteins harboring either the N-terminal (PaGH3.gII.4 and PaGH3.gI.2) or the C-terminal (PaGH3.gII.2 and PaGH3.gI.1) contain the motifs 1 or 3, respectively (**Fig. S5**). The partial protein sequences from PaGH3gII.3, PaGH3gII.5, PaGH3gII.6 and PaGH3gII.10 did not contain any of the 3 motifs and only show conserved residues at the very C-terminal region (**Fig. S5**). The conservation of the 3 motifs in full-length PaGH3 proteins suggests that they might be able to bind ATP/AMP. The *GH3* gene family in Arabidopsis is composed of three subgroups, based on sequence similarities and protein function (Hagen & Guilfoyle, 2002; Okrent & Wildermuth, 2011; Terol *et al.*, 2006). Among members of Group I, GH3.11 displays jasmonic acid (JA)-amino synthetase activity, whereas Group II contains proteins that conjugate IAA and/or JA and SA to amino acids (Staswick *et al.*, 2002; 2005; Terol *et al.*, 2006; Gutierrez *et al.*, 2012; Westfall *et al.*, 2016). An additional cluster including group III was only found in Arabidopsis and some other Eurosids (Terol *et al.*, 2006). To assess the relationships of spruce *GH3* family members to their potential orthologs in Arabidopsis, a phylogenetic tree was constructed including protein sequences from *P. abies* and from Group I and II of Arabidopsis (**Fig. 3**). An additional sequence from *Pinus pinaster* (PpinGH3.16) was included, as it is so far the only *GH3* homologue described in conifers (Reddy *et al.*, 2006). Two out of 14 PaGH3s were found in Group I and thus, were named PaGH3.gI.1 and PaGH3.gI.2 (**Fig. 3**). Interestingly, PaGH3.gI.1 exhibits the same motif variation reported for AtGH3.11 and AtGH3.10 (**Fig. S4**), reinforcing the observed orthology relationship between these GH3 members that cluster together within Group I (**Fig. 3**). Twelve out of 14 PaGH3s were found in Group II (**Fig. 3**). Two of them, such as PaGH3.17 and PaGH3.16, specifically clustered with AtGH3.17 and PpinGH3.16, respectively, and were named according to their potential orthologs in Arabidopsis or *P. pinaster* (**Fig. 3**). The remaining PaGH3s that were found within Group II have no corresponding Arabidopsis orthologs and clustered together in a subgroup closely related to AtGH3.5 and AtGH3.6 and therefore we gave them the generic name of PaGH3.gII (from 1 to 10) (**Fig. 3**). To determine the evolutionary position of the PaGH3 proteins, a phylogenetic analysis was constructed including predicted GH3-like protein sequences from early land plants like liverwort (*Marchantia polymorpha*), mosses (*Physcomitrella patens* and *Sphagnum fallax*) and lycophyte (*Selaginella moellendorffii*) (**Fig. S6**). The split between the two main groups (I and II) occurred very early as moss proteins were present in both main clusters, confirming previous results (Terol *et al.*, 2006) (**Fig. S6**). Most of the GH3-like sequences from *Selaginella* form 2 specific subclusters within the Group I and II, respectively, and only 2 lycophyte proteins are closely related to seed plant main clusters (**Fig. S6**). Within Group II, moss proteins also form a specific subcluster and liverworts are closely related to the lycophyte subgroup. Based on these evidences, the most likely explanation is for the split between the two main groups to pre-date even the most basal land plants and the vascular non-seed plants. Arabidopsis and spruce GH3 proteins have evolved from few copies from that common ancestor, whereas the other land plants (bryophytes and lycophyte) have experienced a separate extension. In seed plants, the topology of the tree reflects the separation of gymnosperms and angiosperms, suggesting that there was a rapid evolution of the GH3 family even before the monocot-dicot split, as indicated by the strict subclustering between Arabidopsis and spruce GH3 proteins (**Fig. S6**).

**Figure 3.**
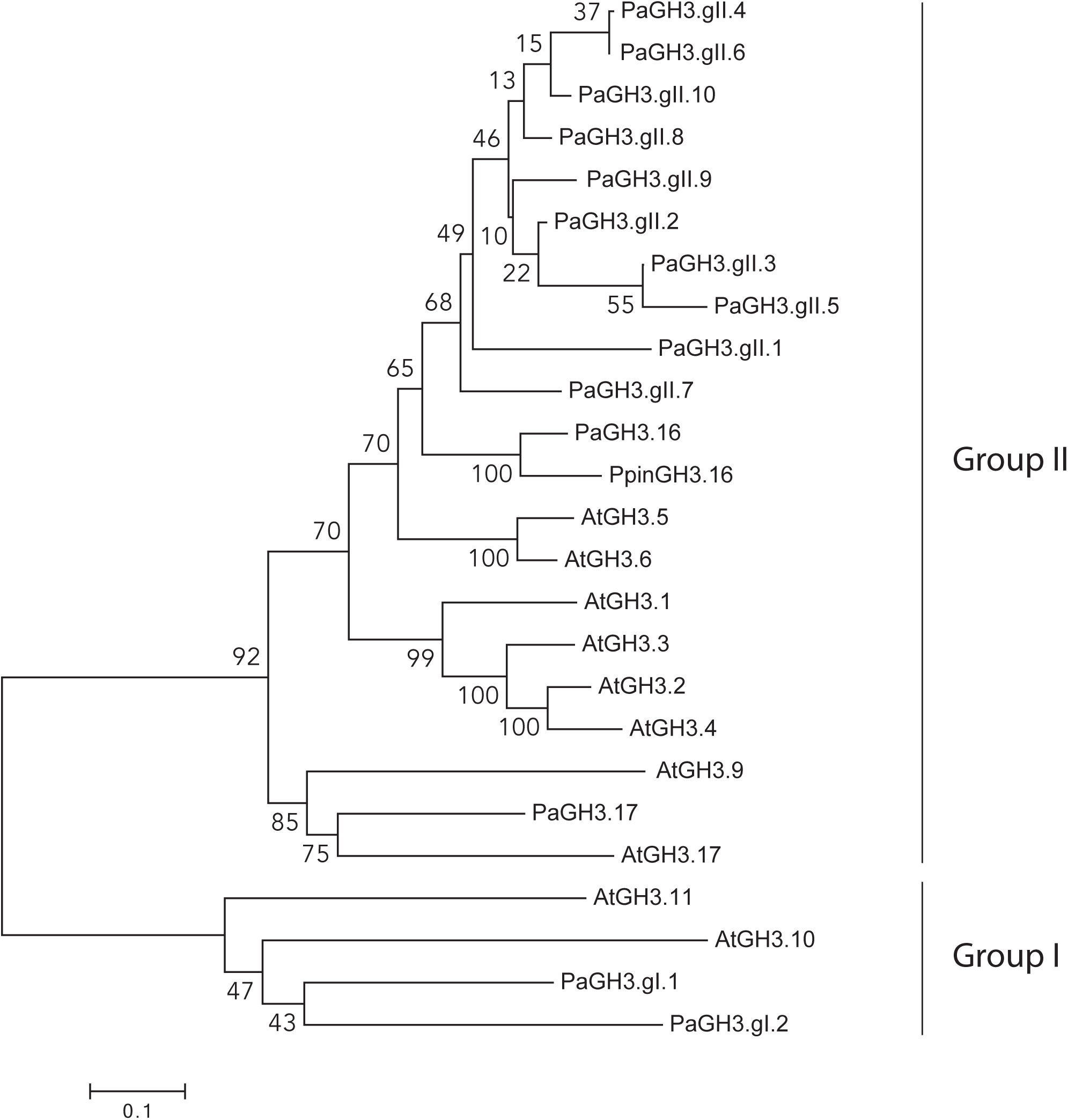
Phylogenetic relationships of GH3 proteins between *Picea abies* and *Arabidopsis*. Predicted proteins sequences (both full-length and partial) from *P. abies* (Pa), *Pinus pinaster* (Ppin) and *Arabidopsis thaliana* (At) were aligned using MUSCLE program. The phylogenetic tree was constructed using MEGA6 program and the Neighbor-Joining method with predicted GH3 proteins. Bootstrap support is indicated at each node.

### 4. A subset of *PaGH3* genes is auxin-inducible and catalyzes the conjugation of IAA to aspartate and glutamate

To determine whether the accumulation of irreversible amide-linked conjugates could be related to modifications of the expression of auxin conjugation genes, the response of *PaGH3* genes to exogenous auxin treatment was analyzed using qRT-PCR. Spruce seedlings were treated with or without 10 μM IAA for 6 hours and gene expression was studied in root tissues. Two *PaGH3* partial genes (*PaGH3.gII.6* and *PaGH3.gII.10*) were excluded from our analysis as the partial nucleotide sequence of *PaGH3.gII.10* was not distinguishable from the full-length sequence of *PaGH3.gII.8* and transcripts from *PaGH3.gII.6* were not detected under our experimental conditions. After auxin treatment, the relative transcript amount of 9 *PaGH3* genes (*PaGH3.gII.9, PaGH3.gII.5, PaGH3.gII.3, PaGH3.*16, *PaGH3.gII.8, PaGH3.gII.2, PaGH3.gII.4, PaGH3.gII.7* and *PaGH3.17*) was increased, while that of *PaGH3.gII.1, PaGH3.gI.1* and *PaGH3.gI.2* was either reduced or not affected (**Fig. 4**). The finding that the 2 *PaGH3* genes from Group I were not induced by auxin is consistent with previous studies which concluded that *AtGH3.10* and *AtGH3.11* were not auxin-inducible genes (Hagen & Guilfoyle, 2002; Winter *et al*., 2007; Okrent & Wildermuth, 2011). This corroborates the observed orthologous relationship of these Arabidopsis and spruce *GH3* genes.

**Figure 4.**
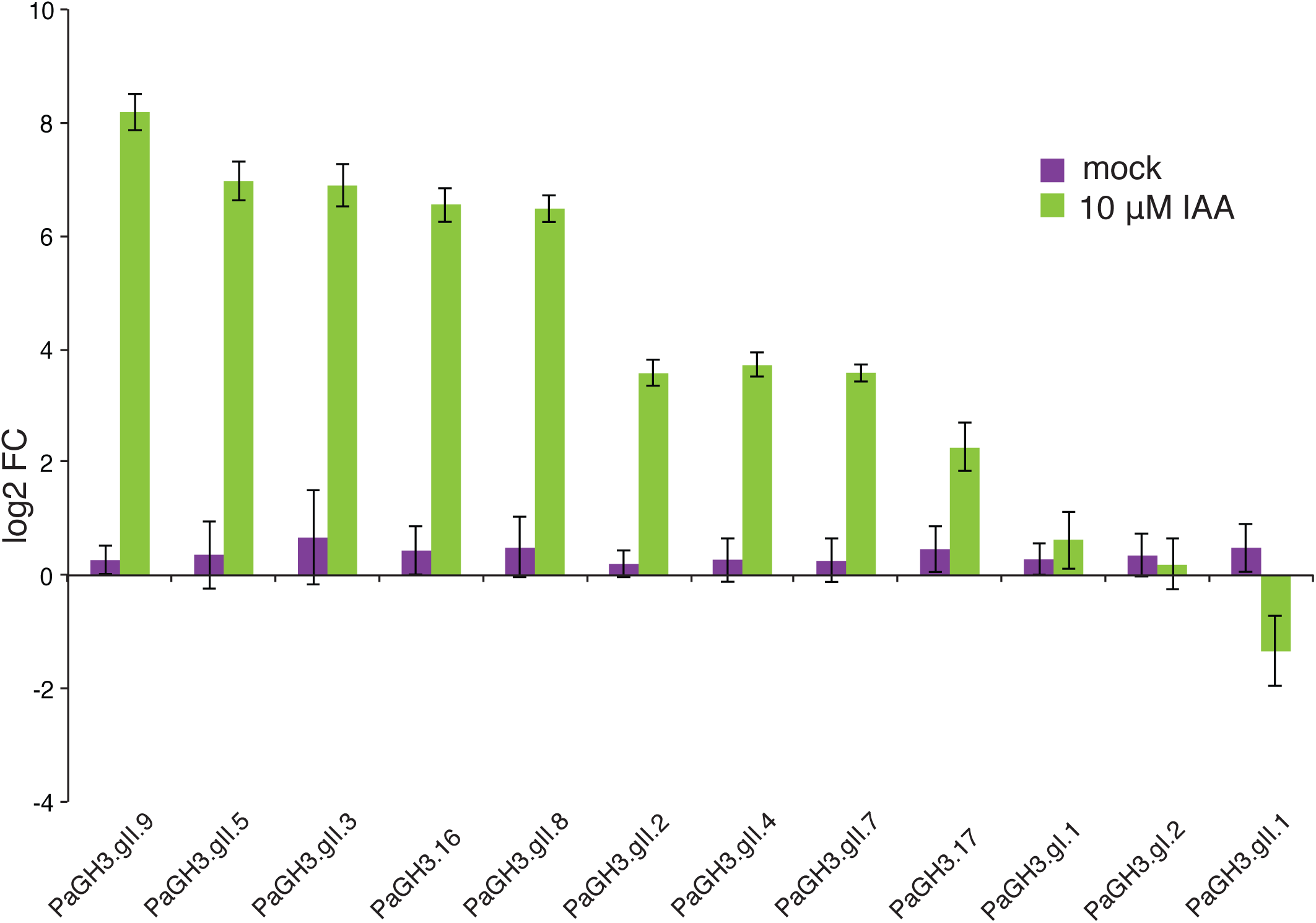
Expression profile of *PaGH3* genes after treatment with auxin. qRT-PCR analysis was performed using cDNA from roots of 2-week-old spruce seedlings treated with or without 10 µM IAA for 6 hours. Fold change (FC) was calculated by comparative Cycle threshold (Ct) method and values were normalized with the expression of the *PaeIF4A* gene. Expression of *PaGH3.gII.6* and *PaGH3.gII.10* genes was not detected in our experimental conditions. Error bars represent ±SD from three biological replicates.

Intrigued by the results from phylogenetic and expression analyses, we cloned 4 of the auxin-inducible *PaGH3* genes and we tested their IAA-conjugating activity. *PaGH3* proteins were expressed in *E. coli* individually, and the expression of recombinant fusion proteins was confirmed by Western blot analysis (**Fig. S7**). Cultures of PaGH3.16-, PaGH3.17-, PaGH3.gII.8- and PaGH3.gII.9-expressing bacteria were tested in a reaction with or without 0.1 mM IAA in combination with GH3 cofactor mixture, as these conditions were previously optimized for AtGH3-mediated IAAsp and IAGlu production (Brunoni *et al.*, 2019). **Fig. 5** shows that, while the reaction mediated by PaGH3.16 led to the accumulation of very little amounts of IAAsp and IAGlu, the reaction mediated by the other PaGH3s tested led to the accumulation of very high levels of IAAsp and/or IAGlu. PaGH3.17 prefered Glu over Asp, similarly to its Arabidopsis ortholog GH3.17 (Staswick *et al.*, 2005; Brunoni *et al.*, 2019) (**Fig. 5**). Interestingly, the reaction mediated by PaGH3.gII.8 yielded the accumulation of both amide-linked conjugates but IAA was conjugated more efficiently with Asp than Glu, as IAAsp levels were much higher than IAGlu (**Fig. 5**). Although to a minor extent compared to PaGH3.gII.8, bacterial cultures expressing PaGH3.gII.9 accumulated higher levels of IAAsp than IAGlu, suggesting that also this enzyme prefers Asp over Glu (**Fig. 5**). Accumulation of IAAsp and IAGlu was always below the limit of detection in all mock samples (**Fig. 5**). Our results demonstrate that at least 3 Norway spruce GH3 proteins, *PaGH3.17, PaGH3.gII.8* and *PaGH3.gII.9*, are able to conjugate IAA with Asp and/or Glu, supporting the finding that these GH3 proteins are responsible for irreversible amide-linked conjugation, thus maintaining IAA homeostasis in spruce.

**Figure 5.**
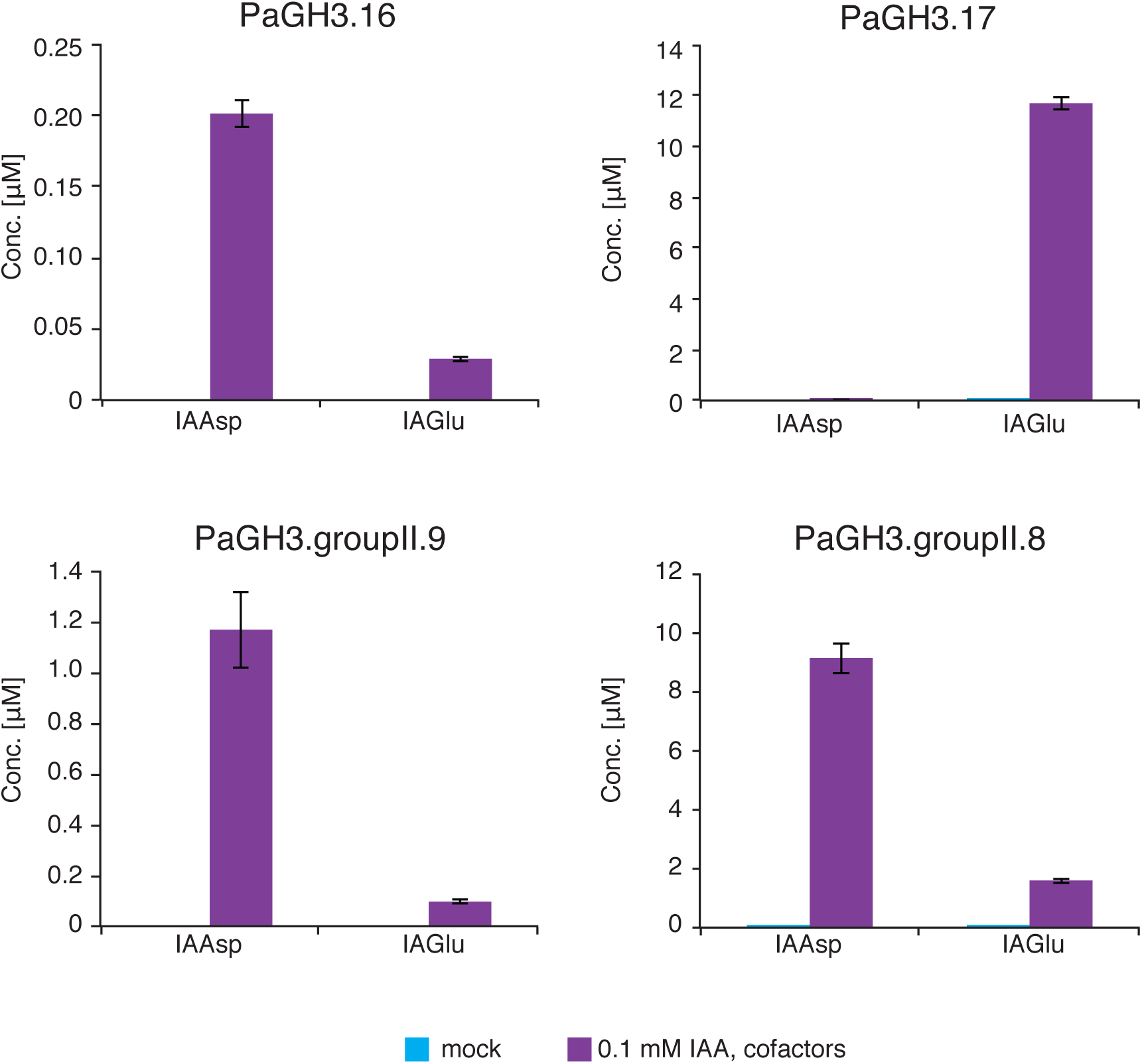
IAA conjugation activity of PaGH3.16, PaGH3.17, PaGH3.gII.8 or PaGH3.gII.9 recombinant proteins. The evaluation of enzyme reactions for IAAsp and IAGlu formation was performed using PaGH3.16-, PaGH3.17-, PaGH3.gII.8- or PaGH3.gII.9-expressing bacterial cultures. All the bacterial cultures were incubated with 0.1 mM IAA and with cofactor mixture (1 mM Glutamic acid, 1 mM Aspartic acid, 3 mM ATP and 3 mM MgCl_2_) for 6 hours at 20°C and only supernatant fractions were analyzed. Bacterial cultures without IAA and cofactor mixture were used as mock. Mean ±SD (*n* = 3).

## Discussion

Maintaining appropriate auxin concentrations is critical for regulating all aspects of plant growth and development. Regulation of auxin homeostasis *via* conjugation and catabolism has been suggested to be a significant contributor to this process. IAA conjugation and degradation pathways and the key components of their machineries have been identified and well characterized in Arabidopsis by incorporating results obtained from gene expression and functionality studies and metabolic analyses. The formation of oxIAA was shown to be major pathway for IAA degradation, acting redundantly and cooperatively with reversible and irreversible IAA conjugation to maintain steady state auxin levels in Arabidopsis. A proposed model for catabolic regulation of IAA in Arabidopsis reveals that the DAO-mediated oxidation pathway finely tunes basal auxin levels under normal growth conditions while the GH3-dependent conjugation pathway responds to both developmental and environmental stimuli (Park *et al.*, 2007; Mellor *et al.*, 2016; Porco *et al.*, 2016).

Our understanding of the physiological mechanisms for modulating IAA levels in gymnosperms is much less understood. Nonetheless, it seems that auxin degradation and conjugation are conserved mechanisms also in gymnosperms, but the contribution of these processes in maintaining IAA homeostasis has not been elucidated yet. Comparative studies of gymnosperms and angiosperms are the key to gain a better understanding of genetic regulation of mechanisms responsible for the diversification of IAA action in seed plants. Within gymnosperms, conifers represent the major division, comprising two-thirds of the extant lineage and they have huge ecological and economic importance. The recent release of genome drafts for spruces and pines have opened new possibilities for studying regulatory gene functions in conifers. In this article, we have studied the contribution of regulatory processes to maintain optimal levels of IAA in seedlings from three different conifer species (*P. abies, P. sylvestris* and *P. contorta*), and investigated the conservation of key molecular mechanisms involved in the degradation of IAA in Norway spruce. The long generation time of conifers makes it difficult to perform studies of gene function, therefore we used young seedlings as an experimental system to study IAA inactivation pathways in conifers. We investigated the shift of IAA metabolite content under steady state conditions and after exogenously applied IAA treatment. Interestingly, we observed that, among the primary IAA catabolites, oxIAA accumulated at very low levels in all the tissues analyzed from conifer seedlings under steady state growth conditions (**Fig. 1; Fig. S2**). On the other hand, we observed that the other two primary catabolites, the amide conjugates IAAsp and IAGlu, contributed more than oxIAA to IAA catabolism in conifers under steady state conditions. Perturbation of IAA homeostasis by feeding with exogenous IAA confirmed the observed metabolic pattern, as conifer seedlings accumulated these two irreversible amide-linked conjugates in much higher concentrations compared to oxIAA (**Fig. 2a; Fig. S3**). A feeding experiment using labeled IAA revealed that IAAsp was the primary IAA catabolite originating from *de novo* synthesis, highlighting the production of IAAsp as the favorite route for IAA degradation in spruce (**Fig. 2b**). In Arabidopsis seedlings, the glucosylester oxIAGlc is the most abundant IAA catabolite (Kai *et al.*, 2007; Porco *et al.*, 2016), and it is more likely synthesized *via* glucosylation of oxIAA and not oxidation of IAGlc, suggesting that levels of oxIAGlc could mirror oxIAA levels. In conifer seedlings, we found that, similarly to oxIAA, oxIAGlc was also detected at very low concentration, confirming that the oxidation pathway plays a minor role in modulating IAA levels in conifer seedlings compared to Arabidopsis (**Fig. 1; Fig. S2**).

In conifers, glucose conjugation to IAA also contributed to the inactivation of IAA under steady state conditions or after exogenous treatment with IAA (**Fig. 1; Fig. S2; Fig. 2a; Fig. S3**). The formation of IAGlc is a well-known mechanism for IAA inactivation and this ester-linked conjugate is considered as a reversible storage form, which level depends on the conjugation/hydrolysis rates (Jackson *et al.*, 2001).

Together, these findings indicated that, on one hand, oxIAA formation contributed the less to the maintenance of steady state IAA levels in conifer seedlings, both under steady state growth conditions and after treatment with exogenous IAA, and, on the other hand, the conversion of IAA to IAAsp and IAGlu appeared to be a major catabolic pathway, acting at a constitutive level and after IAA homeostasis perturbation in conifer seedlings. These findings do not preclude that these mechanisms could differentially contribute to reduce IAA levels at other developmental stages or under different growth conditions and that conifers could possess other IAA inactivation pathway(s) that have not been identified yet.

The GH3 auxin-conjugating enzymes from Group II mediate the formation of IAAsp and IAGlu in Arabidopsis and this pathway seems to be highly conserved all over the plant kingdom (Staswick *et al.*, 2005; Terol *et al*., 2006; Reddy *et al.*, 2006; Ludwig-Müller *et al.*, 2008; Ludwig-Müller, 2011). We have therefore focused on the molecular identification of spruce GH3 homologues that could be involved in the formation of the amide-linked conjugates IAAsp and IAGlu. Phylogenetic analysis showed a homology clustering consistent with the functional classification of the Arabidopsis proteins, since the PaGH3 sequences grouped into the 2 main clusters, corresponding to the Arabidopsis functional group I (JA adenylation) and II (IAA adenylation) as shown in **Fig. 3**. A comparative phylogenetic analysis including predicted GH3-like protein sequences from early land plants confirmed that the GH3 family appeared very early in the plant kingdom and differentiated into the two main groups (I and II) very soon (**Fig. S6**), confirming previous findings reported by Terol *et al.* (2006). In seed plants, the topology of the tree reflects the gymno-angiosperm split, as indicated by the strict subclustering between Arabidopsis and spruce GH3 proteins, suggesting that the evolution of the family pre-dated the separation of gymnosperms and angiosperms (**Fig. S6**). In Arabidopsis, the expression of several Arabidopsis *GH3* genes of Group II is induced by exogenously applied auxin, while *AtGH3.9, AtGH3.17* and the *AtGH3s* of Group I show little or no induction by auxin (Hagen & Guilfoyle, 2002; Winter *et al*., 2007; Okrent & Wildermuth, 2011; Di Mambro *et al.*, 2017). *PpinGH3.16*, the only other Group II *GH3* homologue from conifers studied so far, was also specifically up-regulated after auxin treatment (Reddy *et al.*, 2006). It has been suggested that plants have developed a feedback mechanism to control the levels of active auxin through modulation of *GH3* gene expression by auxin itself (Staswick *et al.*, 2005). Expression analysis of *PaGH3* genes revealed that 9 *PaGH3s* of Group II were highly up-regulated after treatment with IAA, while the other *PaGH3* of Group II (*PaGH3.gII.1*), and *PaGH3s* of Group I were either downregulated or not induced by auxin (**Fig. 4**). The differential response to auxin among the *PaGH3* genes mirrors the phylogenetic clustering of the corresponding enzymes (**Fig. 3**). Most of Arabidopsis Group II enzymes are demonstrated to be active on IAA and each GH3 enzyme has a broad specificity for amino acids and could synthesize several types of IAA-amino acid conjugates (Staswick *et al.*, 2005). Nonetheless, evaluation of GH3 enzyme activity with different amino acids *in vitro* and GH3 multiple mutant analyses revealed that IAAsp is a major conjugate formed by AtGH3.1-6 and that IAGlu formation is mediated by AtGH3.17 (Staswick *et al.*, 2005; Porco *et al.*, 2016). Heterogously-expressed AtGH3.6 and AtGH3.17 also efficiently conjugate IAA with Asp and/or Glu in a bacterial assay (Brunoni *et al.*, 2019). We have therefore investigated possible IAA-conjugating activity with Asp and Glu for 4 of the auxin-inducible PaGH3-recombinant proteins by performing the enzymatic assay directly in bacterial cultures (**Fig. 5**). Among them, PaGH3.17 enzyme conjugated IAA mainly to Glu, supporting the assumption that PaGH3.17 and AtGH3.17 could be functionally equivalent orthologous. On the other hand, PaGH3.gII.9 and PaGH3.gII.8 appeared to favor Asp over Glu, as IAAsp accumulated more than IAGlu. Here, we have found that these GH3 proteins are responsible for irreversible amide-linked conjugates and thus, could function redundantly in reducing IAA concentrations in spruce.

Considerable evidence has been accumulating in support of the notion that the pathways for IAA oxidation and conjugation are the predominant IAA homeostatic pathways operating in all the land plants and algae. Cooke *et al.* (2002) proposed that while both charophytes and liverworts use a more primitive regulatory strategy to deal with excess of IAA, which is based on the control of the balance between the rates of IAA biosynthesis and IAA degradation, more sophisticated machineries have evolved afterwards. As a result, IAA conjugation and conjugate hydrolysis reactions have developed and started to play a more important role in the regulation of auxin homeostasis in all the other land plants. It is believed that this metabolic implementation allows plants to achieve a more precise spatial and temporal control of IAA accumulation (Cooke *et al.*, 2002). Since then, new methods for more precise and sensitive IAA metabolite profiling were developed and used to retrieve data from several species ranging from cyanobacteria and algae to several bryophytes (Drábková *et al.*, 2015; Žižková *et al.*, 2017). The bulk of new data available highlighted that oxidation is a major pathway for IAA degradation in algae, in basal land plants and angiosperms, while it seems to be of minor importance in conifers (**Fig. 6**). Arabidopsis have developed an IAA homeostatic response based on a slow oxidative mechanism that modulates constitutively the optimal auxin levels and a fast conjugative and inducible pathway that reduces IAA from the active auxin pool upon stress or developmental cues (**Fig. 6**). This metabolic pattern seems to arise from pre-existing elements of IAA inactivation machinery already operating in green algae and mosses and contributing to regulate levels of active auxin similarly to Arabidopsis. First of all, IAA catabolism *via* oxidation was found to be a more relevant homeostatic mechanism than the conjugative pathway in cyanobacteria, algae and the most basal land plants (**Fig. 6**) (Drábková *et al.*, 2015; Žižková *et al.*, 2017). Secondly, results from feeding several algae species with IAA demonstrated that the IAA conjugative pathway is rapidly activated, as IAAsp and IAGlc were the two major products. Interestingly, the metabolization of exogenous labeled IAA in these selected algal species led to the detection of several more unidentified metabolites, indicating the existence of other unknown IAA metabolic pathway(s) in green algae (Žižková *et al.*, 2017). On the other hand, conifers seem to prefer to rely mainly on a conjugative mechanism for fine-tuning basal IAA levels under steady state conditions and that can be rapidly activated in response to exogenously applied IAA (**Fig. 6**). Taken together our findings suggest that the strategy adopted by conifers to maintain homeostatic level of endogenous IAA differs from all other land plants and algae studied so far. Further studies in other gymnosperms taxa and at other developmental stages would add further tiles to confirm that this diversification of IAA action is conserved in the lineage of gymnosperms.

**Figure 6.**
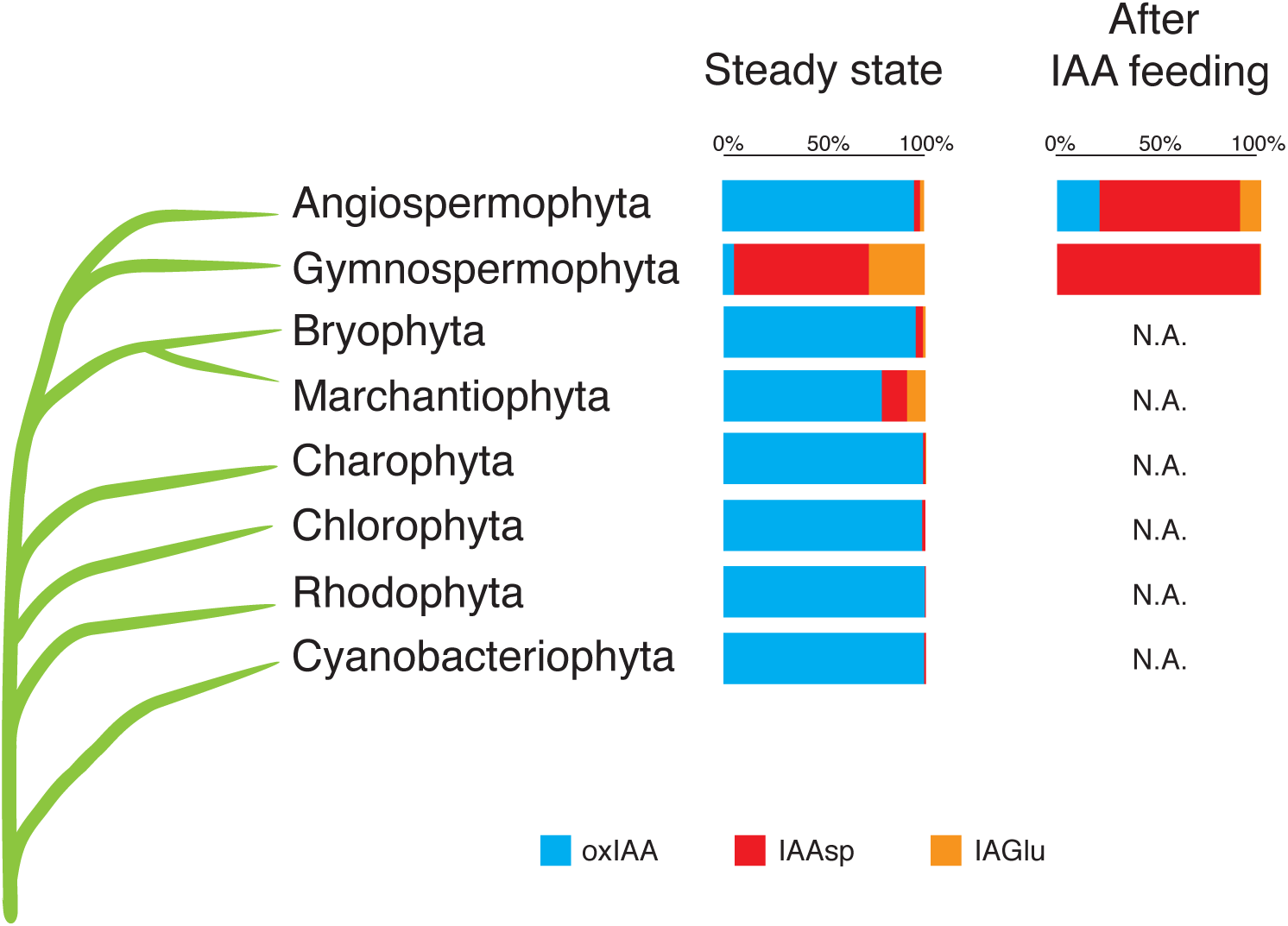
Schematic illustration of the most studied IAA inactivation pathways in land plants, algae and cyanobacteria. Proportion of oxIAA, IAAsp and IAGlu in the total IAA catabolite pool under steady state conditions for Cyanobacteria, Rhodophyta, Chlorophyta, Charophyta (Žižková *et al.*, 2017), Marchantiophyta, Bryophyta (Drábková *et al.*, 2015), Gymnospermophyta (this study) and Angiospermophyta (Porco *et al.*, 2016) is reported. Proportion of IAA irreversible catabolites after feeding with labeled IAA for angiosperms (Porco *et al.*, 2016) and gymnosperms (this study) is reported.

## Acknowledgements

This research was supported by grants from the Swedish research councils FORMAS, VR, Kempestiftelserna, and the Knut and Alice Wallenberg Foundation (to KL and CB). The authors would like to thank Dr. Nicolas Delhomme for helpful discussions and the Swedish Metabolomics Center (SMC, Umeå, Sweden) for access to instrumentation.

## Authors’ contributions

FB, SC, KL and CB conceived and designed the experiments. FB, SC, RCS, JS and MK performed all the experiments. FB and SC wrote the manuscript with the input from all authors. All authors read and approved the final article for publication.

