## Supporting information for "Conifers exhibit a characteristic inactivation of auxin to maintain tissue homeostasis"

**The following Supporting Information is available for this article:**

Table S1
Fig. S1 to S7

Table S1: Primers used in this work.

| Purpose | Name | Gene ID | Primer sequence (5'→3') Forward | Reverse |
|---|---|---|---|---|
| RT-qPCR | PaGH3.16_q | MA_10434772g0010 | GCTACCACCGAGCGTAACAAG | CACTGAACCCATAGCGTTGAAG |
| | PaGH3.17_q | MA_16777g0010 | CGGTCGAATCGATAACAAATCG | CTGTGAAGCTCCTCGGGAAT |
| | PaGH3.gII.1_q | MA_100975g0010 | CACGAGGTGGTATAAGAGTCG | GAACCCAGAAGCAAAGATGGC |
| | PaGH3.gII.2_q | MA_10432413g0010 | CACTGGATGAGAAAGAGCGAC | GCGTCGATACTCAGCACAA |
| | PaGH3.gII.3_q | MA_580596g0010 | CTATCTATCGCCAAGGAAGAGC | GGAGTTTAGCAGCTCAACGA |
| | PaGH3.gII.4_q | MA_66253g0010 | GACGATCTCAAGTCTGAG | CGGGCATGAGAAGACTATACAG |
| | PaGH3.gII.5_q | MA_6467862g0010 | GGTCACTACGTGCTATACTG | CCCTCTACTGAGAGCATAATCC |
| | PaGH3.gII.7_q | MA_212507g0010 | CGTCGTATTGCTAACGGAGAC | CCCTTGTCAAGTCCTTGCA |
| | PaGH3.gII.8_q | MA_10432413g0020 | CTGCTCATCCAGTTTCCGAA | GGAGTCTTTGCCTCCGATT |
| | PaGH3.gII.9_q | MA_10330250g0010 | CGAGAATACGAGCTTGTCG | CAGCTCTGCTTCATCCGTTT |
| | PaGH3.gI.1_q | MA_10429520g0010 | CATCTGCGTCATGAAGGCTT | CGGTAACTTTCACGACATCACC |
| | PaGH3.gI.2_q | MA_376158g0010 | GACTGGAGACGGAAACCAGAA | CCAAGTCCACTTGCATTGC |
| | PaeIF4A_q | MA_50378g0010 | TTGGTCGGAGTGGACGATTTGG | TGACGAGAGAATGCTGCAGGAC |
| Cloning | PaGH3.16_clo | MA_10434772g0010 | TCCCCTCTAGAAATAATTTTGATTTAACTTTAAGAAGGAGATATACCATGCATCACCATCACCATCACGCGATGGGAGAGAAAGCGAAAGC | CTCGAGTGCGGCCGCAGCAAGCTTCTAGCTAGCTTATTATTACCTCACAATTTTAGCAACAAAC |
| | PaGH3.17_clo | MA_16777g0010 | TCCCCTCTAGAAATAATTTTGATTTAACTTTAAGAAGGAGATATACCATGCATCACCATCACCATCACGCGATGGAGATGAATATTCGCTGC | CTCGAGTGCGGCCGCAGCAAGCTTCTAGCTAGCTTATTATTAATAAAGCTGAGAATTTGG |
| | PaGH3.gII.8_clo | MA_10432413g0020 | TCCCCTCTAGAAATAATTTTGATTTAACTTTAAGAAGGAGATATACCATGCATCACCATCACCATCACGCGATGAGTTCTTCTCTGGGAAACGAGG | CTCGAGTGCGGCCGCAGCAAGCTTCTAGCTAGCTTATTATTATGCCCAATTTCTGCATCCTGG |
| | PaGH3.gII.9_clo | MA_10330250g0010 | TCCCCTCTAGAAATAATTTTGATTTAACTTTAAGAAGGAGATATACCATGCATCACCATCACCATCACGCGATGAGTTCTGCTATGGGAAAC | CTCGAGTGCGGCCGCAGCAAGCTTCTAGCTAGCTTATTATTACGCCCATTCTCTGCATCC |
| 5'-RACE | PaGH3.gII.9_gsp | MA_10330250g0010 | | TGAGCCATTGCACCAGTGACGATGACAT |

**Fig. S1**

**Full-length coding sequence of *PaGH3.16*, *PaGH3.17*, *PaGH3.gII.8* and *PaGH3.gII.9*.**

```
>PaGH3.16_CDS

ATGCCTCAGGCAAAGAGAGAAGAGAGTATGGAGTCCGTCCAAGAGTGTATTAGCATTGCTACCACCGAGC
GTAACAAGGAAGCTCTGGATTTCATCGAGAATGTCACCGTTCATGCTGATGCAGTGCAAGAGCGAGTTCT
GTTGGAAATTTTAACTAGAAATGCTCATACAGAATATCTTCAACGCTATGGGTTCAGTGGTCGCACTGAT
AGAGAGTCATTCAAGGAATGTTTTCCTGTGATTACTTACGAAGATCTGCAGCCTGAGATTCTTCGCATAG
CTAACGGAGACACATCTCCCATTCTTTCTGCTCACCCCATCTCTGAATTTCTCACAAGCTCTGGAACATC
GGCTGGAGAGCGCAAGATTATGCCCACTATTCACGAAGAATTAGAGCGGAGAACGCTTCTTTACAGCCTG
CTTATGCCTGTCATGAACCAATACATGGAAGGGCTGGACAAGGGAAAAGGCATGTACTTCCTCTTCATCA
AATCTGAGACCAGGACTCCCGGTGGGTTGCTCGCTCGTCCTGTCCTCACGAGCTACTACAAGAGTCAGCA
CTTCAGAGAGCGGCCTTACGACCCCTACAATGTATACACCAGCCCCATAGAGGCGATACTAAGTGCAGAC
TCATATCAAAGCATGTACTGCCAGCTTCTCTGCGGTCTTGCGCAGAACCACGAGGTGCTCAGAGTGGGAG
CCGTGTTCGCGTCCGGGCTTCTGAGAGCAATACGTTTCCTGGAGGAACATTGGAAGTCGCTGTGTCAAGA
TATTCGCAGTGGCACAATAAACGATGAAGAGGTGACGGATCCATGCCTCAGAGAGAGCGTCATGAAGATA
CTCCGCCCGAAAATTCAACTGGCTGACTTGATCCACGCCGAGTGCTCCAAAGAATCATGGCAAGGAATCA
TCACTCGCCTGTGGCCCAACGCCAGATATTTGGACGTCATTGTTACAGGCGCCATGGCGCAATACATCGA
GACGCTGGATTTTTACAGCGGCGGCCTTCCTCAGGTTTGCACAATGTATGCCTCCTCGGAGTGCTATTTC
GGAATCAATCTCAAGCCTCTGTGCAAGTCGTCCGAGGTTTCCTACACGCTAATGCCCAACATGGCCTTTT
TCGAGTTCCTTCCGGTCTACCGTAACAACGACGATGCAGCGCCCGTCACCATGGCCACCGAGCAGCAAGA
GCTTGTCGATCTAGCTGATGTGACAGTGGGACAAGAATATGAGCTTGTGATCACCACATATGCAGGACTA
TACCGTTACAGAGTGGGAGACGTGCTACGGGTGACGGGGTTCTACAACGCGGCGCCGCAGTTCCAGTTCG
TGTGCAGGAAGAATGTAATGCTGAGCATCGACTCCGATAAAACCGACGAAGCAGAGCTGCACAGCGCAGT
AAAGAATGCTGTTAAGCATCTGGAGCCCTTCGAGGCTAGCCTCGTGGAGCACACCAGCTACGCCGAAACG
TCCACCATTCCGGGTCATTACGTTCTTTACTGGGAGCTGCGGAACTCCACGGTGCCCGTGCCTGCCTCCG
TTTTTGAAGATTGCTGCCTGACCATAGAAGAATCTCTCAACTCCGTCTACAGGCAGTGCAGAGTAGCAGA
CAAATCCATCGGCCCCTTGGAGATTAAAGTAGTTGAAACGGGAACATTCGACAAATTGATGGACTATGCT
ATCAGCCGAGGTTCCTCCATCAATCAATACAAAGCGCCACGATGCGTCAAGTTCGCTCCAATGGTGGATC
TCCTAAAGTCCAGAGTTTCGGCAAGCTATTTCAGCCCCAGGTGTCCCAAATGGACACCAGGACGGACCCA
GTGGGGCGCCATTACTCGGTTAATTTAATAATAA
```

>PaGH3.17_CDS

```
ATGGAGATGAATATTCGCTGCGCCAAAGACAAAGGCGAGGCGGCCCTGCAGCTTATCGAGAATTTGACTG
CCAGGGCCGACGAGGTGCAGAAGCAAGTATTGTATGAGATCTTAAGGAGAAACGCAGAGACCGAGTACCT
GAACAAATTTCTCAACGGTCGAATCGATAACAAATCGTTCAAAATCAACGTACCCGTTGTTAACTACGAG
GATATCAAACCATACATCCAGCGGATAGCCAATGGAGACGCTTCTGCCATTATTTCTGCTGAACCCATAT
CAGAGCTGCTGACAAGCTCTGGAACATCAGGCGGACAGCCTAAAATAATGCCTTCCATTCCCGAGGAGCT
TCACAGGAAAACCTTTCTTTACAATCTACTGATGCCAATAATGAACAAGTACAGGAAAACCTTTCTTTAC
AATCTACTGATGCCAATAATGAACAAGTATGTTCCTGGACTGGACAAAGGAAAAGGGATGTACCTCCAGT
TTATAAAAACGGAGGTAACTACTCCTTCCGGACTAAAGGCGAGGCCGGTGCTGACGAGCTACTACAAGAG
TAGCAACTTCAGGGATCGACCATTTGATAAGTTTAATGTGTATACGAGCCCGGACGAGACAATCCTTTGT
CCCGACAGCAGACAAAGTATGTTTTGCCAATTACTGTGTGGATTATTACAAAGGGACGAAGTACTCAGAG
TTGGGGCTGTTTTTGCCTCCGCATTTCTCAGGGCCATTAAGTTTCTGGAGGAAAACTGGGAAGAACTTTG
CGACAACATACGCACAGGGCATCTGAGCGACTGGATCGACGATCCTCCATCCAGAATTGCTGTTATGAAG
ATGCTCAGTCCTAACCCCCAACTCGCAGAAGAAATCCATGGGGAATGCAGCAAGAAATCGTGGCAAGGAA
TAATTACCCGTCTGTGGAGAAAGACAATATACATAGATGTAATAGTCACTGGAACCATGGCACAGTATAT
TCCAACTCTTGATTACTATGGTGGAGGGCTCCCTTTAGTATCTACCATGTACGCGTCTTCTGAGTGTTAT
TTCGGAGTAAATCTGAAACCTTTGAAAGTCTTAGATTTGCAGTCCAACGGCACAGAAAATGGGAAAAAAA
TAGAGGATGAATTAGTGGACCTGGTGGATGTGAAGGTTGGTCACTACTATGAACTTGTAGTGACAACTTA
TGCTGGCTTATATCGATATAGGGTCGGTGACATTCTCTTAGTGACAGGTTTCTACAACAGAGCCCCTCAG
TTTGAATTTGTTTACCGTAGAAACGTGGTTCTCAGTATTGACACAGACAAAACAAATGAAGAAGATTTGT
TGAAGGCTGTCACTAAAGCCAAGAAATTGTTAGAGCCATTCAATGCCCTGTTATCGGAGTACACTAGCTA
TGCTGATACATCTACACTACCCGGTCACTATGTTCTGTTCTGGGAACTGAATACCAGGGAAGAATTTTTG
GATGCATCCGTATTGGAAAGCTGTTGCTCAACGATCGAGGAATCCTTGGATTCTATCTACAGAAGATGCA
GAACCAAAGATAAGTCTATTGGCCCGCTAGAGATAAGGCTAGTGAAACCTGGGACGTTTGATTTGTTGAT
GGATTATTGTCTGAATCAAGGCTCTTCCTACAATCAGTACAAAACACCCAGATGTATCAAGTCTCTCCAT
GTCCTAGGGCTTCTAAATTCAAAAGTCACTGCCAAATACTTCAGCAAACGTCTTCCGTCTTGGACACCAT
ACAACCCTGGCAGCCTGAATCCAAATTCTCAGCTTTATTAA
```

>PaGH3.gII.8_CDS

ATGAGTTCTTCTCTGGGAAACGAGGTCGATGATCGAAACAAGAGAGCTCTGGAATTCATCGAGACCGTCA
CCACAGACGCAGACGAGGTGCAGACTCAGGTCCTCTCCTCAATCTTATCGAGAAATGCAGACACGGAATA
CTTGAAGAGATACGGCCTCAATGGACGAACTGATAGAGCCACTTTCAAAAAGTGTCTGCCAGTAATCACT
TATGATGATCTGAAGCCTGAAATACGTCGGATTGCTAGCGGAGACACTTCTCCAATCCTTTCTGCTCATC
CAGTTTCCGAATTCCTCACAAGCTCGGGAACCTCTGCTGGTGAACGGAAGCTGATGCCCACCATCCAGGA
GGAATTGGAGAGGAAAGCTCTTCTGTATAGTCTTCTCATGCCCGTCATGAATCAATACATGAAAGGACTT
GACGAGGGAAAGGGAATGTATTTCTTCTTCATAAAATCGGAGGCAAAGACTCCAGGAGGGTTGCTCGCAC
GTCCCGTCCTGACGAGCTATTACAAGAGCGACTACTTCAAGGAGAGGCCTTACGATCCCTACAATGTTTA
CACTAGTCCTAATCAGACAGTCCTCTGTCAGGATGCCTACCAGAGCATGTACTCTCAACTGCTATGCGGT
CTCCTTCAAAATAACGAGGTTCTAAGAATGGGAGCCGTCTTCGCCTCTGGATTCATCAGAGCTATACGCT
TCCTAGAAGAGCACTGGAGACAATTTTGCCTGGACATAAAAACCGGCATTCTCAACAGAGAGGTGACGGA
TCCATCAGTGAGGGAGGCCGTCGGGGAGCTTCTGCACCCGAATCCAGAGTTGGCTGATTTTGTTGAAAGG
AAATGTTCAGCCCAGTCCTGGCAAGGAATTATAACTCGCCTGTGGCCTAACACTAAGTATATCGATGTCA
TCGTGACTGGTGCAATGGCTCAATATATTCCTACTTTGGATTACTACAGCGGCGGGCTACCCCTGGTATG
CACCATGTACGCTTCATCCGAATGCTACTTTGGACTTAATCTGAAGCCTCTCTGCAAGCCTTCAGAAGTG
TCATACACTTTACTCCCAAACATGGCCTATTTCGAGTTCCTTCCCGTTCACCGCAAACAAGAAGCAGCTG
GTCTGACCATAGAATCGTCAACGATCCCCAAAACACTCGACGACAAAGAGCGAGAGGAATTGGTTGAGCT
TGTGGACCTGAAACTTGGGCAAGAATACGAGCTCGTCGTCACCACATATGCTGGTTTGAATCGTTATAGA
GTGGGGGATGTGCTACGCGTGACTGGTTTCCACAACGCTGCGCCTCAATTCCATTTTGTATGCAGACAGA
ATGTTGTTCTGAGTATAGACTCAGACAAAACAGATGAAGTAGAGCTTCACAGTGCGGTAGAGAATTCTGT
CAAGCATCTGGAAGCGTTCGACGCCCAATTGATCGAGTATACAAGCTATGCCGACACAGGTACAATCCCT
GGCCACTACGTGCTATACTGGGAACTTCGCTTCAACACCAAAGCAATTGAAGTTCCTTCTTCAGTTTTTG
AAGATTGCTGTCTCACCGCAGAAGAATCCCTGAATTCTGTCTATCGCCAAGGAAGAGCGTCCGATAAGTC
CATCGGCCCTCTGGAAATTAGGGTAGTTGAAGAGGGAACGTTCAACGAATTGATGGATTATGCTCTCAGT
AGAGGGGCATCCATAAATCAATACAAAGCCCCCAGATGCATTAAATTCACTCCCATCGTTGAGCTGCTAA
ACTCCAGGGTCGTGCATTCCTACTTCAGCCCGCAGCCTCCACAATGGGCTCCAGGATGCAGAAATTGGGC
ATAA

>PaGH3.grII.9_CDS

ATGAGTTCTGCTATGGGAAACGGCGTCAGTGATCGAGACAAGAGAGCTCTGGATTTCATCGAGAGCGTCA
CCACAAACGCAGACGATGTGCAAATTCGTGTCCTCTCCTCGATCTTATCGAGAAATGCGGACACCGAATA
CCTCAAGAGATACGGCCTGAATGGACGAACTGATAGAGCCACTTTCAAAGAGTGTCTGCCAGTGATAACT
TATGACGATCTCCAGCCTGATATACGTCGTATTGTTGGCAGAGACACTTCTCCCATCCTTTCTGCTCACC
CAGTTTCTGAATTCCTCACTAGCTCGGGAACCTCGGCTGGTGAACGGAAGCTGATGCCCACCATCCAGGA
GGAATTGGAGAGGAAAGCTCTTCTGTATAGTCTTCTCATGCCCGTCATGAATCAATATATGCAAGGACTT
GACAAGGGAAAGGGAATGTATTTCTTCTTCATAAAATCGGAGGCAAAGACTCCCGGAGGGTTGCCCGCAC
GTCCCGTCTTGACGAGCTATTATAAAAGCCCCCACTTCAGGGAGAGGCCCTACGATCCCTACAACGTTTA
CACCAGTCCTAACCAGACGGTCCTCTGTCCGGATGCCTACCAGAGCATGTACTCTCAACTGCTATGCGGT
CTACTCCAACATAATGAGGTGCTGAGGATGGGAGCCGTCTTCGCTTCTGGGTTCATCAGAGCTATACGCT
TCCTGGAAGAGCACTGGACACAATTTTGCCTGGACATAAGAACCGGCATTATCAACAGAGAGGTGACGGA
TCCATTAGTGAGAGAGGCCGTCGGGGAGCTTCTGCACCCGAATCCAGAGTTGGCCGATTTGGTTGAAAGG
GAATGTTCAGCCCCGTCATGGCAAGGAATTATAGCTCGCCTCTGGTCTAACACTAAGTATATCGATGTCA
TCGTCACTGGTGCAATGGCTCAATATATTCCTACTTTGGATTACTACAGTGGCGGGCTACCACTGGTATG
CACCATGTACGCTTCGTCCGAATGCTACTTTGGACTTAATCTGAAGCCTCTCTGCAAGCCTTTTGAAGTG
TCATACACTTTACTCCCAAACATGGCCTATTTTGAGTTCCTTCCCGTTCACCGCAAACAAGAAGAAGCTG
GCGTGACTCTCGAATCGCCAGGGACACTCGACGAGAAAGAGCGAGAGGAGTTGGTGGAACTTGCTGACGT
GAAACTTGGGCGAGAATACGAGCTTGTCGTCACCACGTATGCGGGTTTGAATCGTTATAGAGTCGGGGAT
GTGCTGCGTGTGACTGGATTTTACAACGCTGCGCCACAATTCCATTTCGTGTGTAGACAGAATGTTGTGC
TCAGCATCGACTCCGACAAAACGGATGAAGCAGAGCTGCACAGCGCGGTAGAGAATTCTGCTAAGCATCT
GGTGCCATTCCACGCCCAATTGGTCGAGTATACAAGCTATGCCGACACAGCGACAATACCTGGTCACTAC
GTGCTGTATTGGGAACTTCGCTTCGACACCAAAGCAGTTGCATTTCCTTCTTCAGTCTTTGAAGATTGTT
GTCTCACTATAGAAGAATCCCTGAATTCTGTTTATCGCCAAGGAAGAGCGTCCGACAGGTCCATCGGCCC
CCTGGAAATTAGGGTAGTTGAAGAGGGAACGTTCGACCAATTGATGGATTATGCTCTCAGTAGAGGGGCA
TCCATAAATCAATACAAAGCTCCCAGATGCGTTAAATTCACACCCATTGTTGAGCTGCTAAACTCCAGGG
TCGTGCATTCCTACTTCAGCACCCAGCCACCACAATGGGCTCCCGGATGCAGAGAATGGGCGTAA

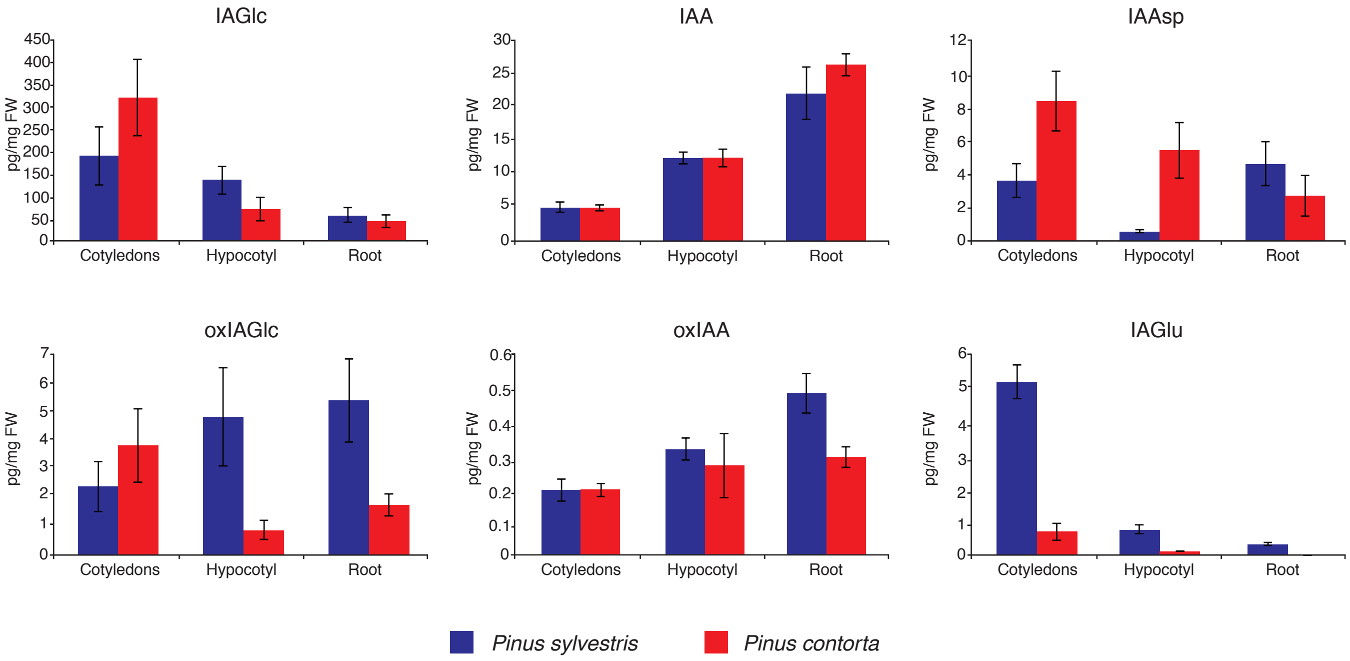


**Figure S2**
**Levels of IAA metabolites in different organs of *Pinus sylvetris* and *Pinus contorta* seedlings.**
IAA and IAA metabolites oxIAA, oxIAGlc, IAGlc, IAAsp and IAGlu were quantified in cotyledons, hypocotyl and root from 2-week-old pine seedlings. The level of IAGlu in *P. contorta* root was under the detection limit of the used LC-MS/MS method. Blue indicates *P. sylvestris* and red indicates *P. contorta*. The concentrations of all metabolites are in picograms per milligram fresh weight (FW). Error bar indicates ±SD (n = 4).

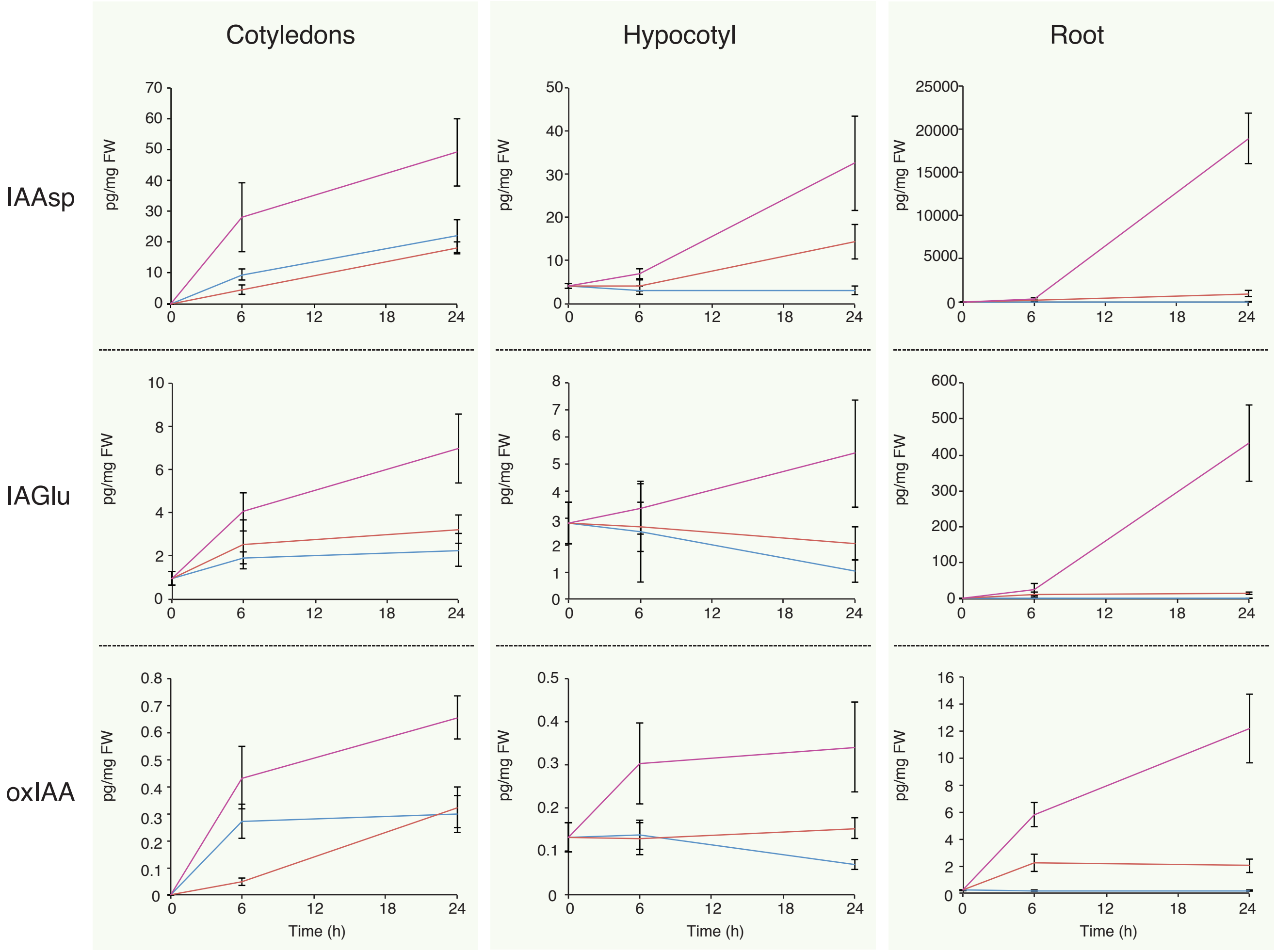


**Figure S3**
**Concentrations of IAA metabolites in different organs of *Picea abies* after feeding with unlabeled IAA.**
Two-week-old spruce seedlings were incubated with 1 (red line), 10 μM (pink line) or without (blue line) unlabeled IAA and the levels of IAA and IAA metabolites IAAsp, IAGlu and oxIAA were quantified in cotyledons, hypocotyl and root after different incubation times. The concentrations of all metabolites are in picograms per milligram fresh weight (FW). Error bars indicate ±SD (n = 4).

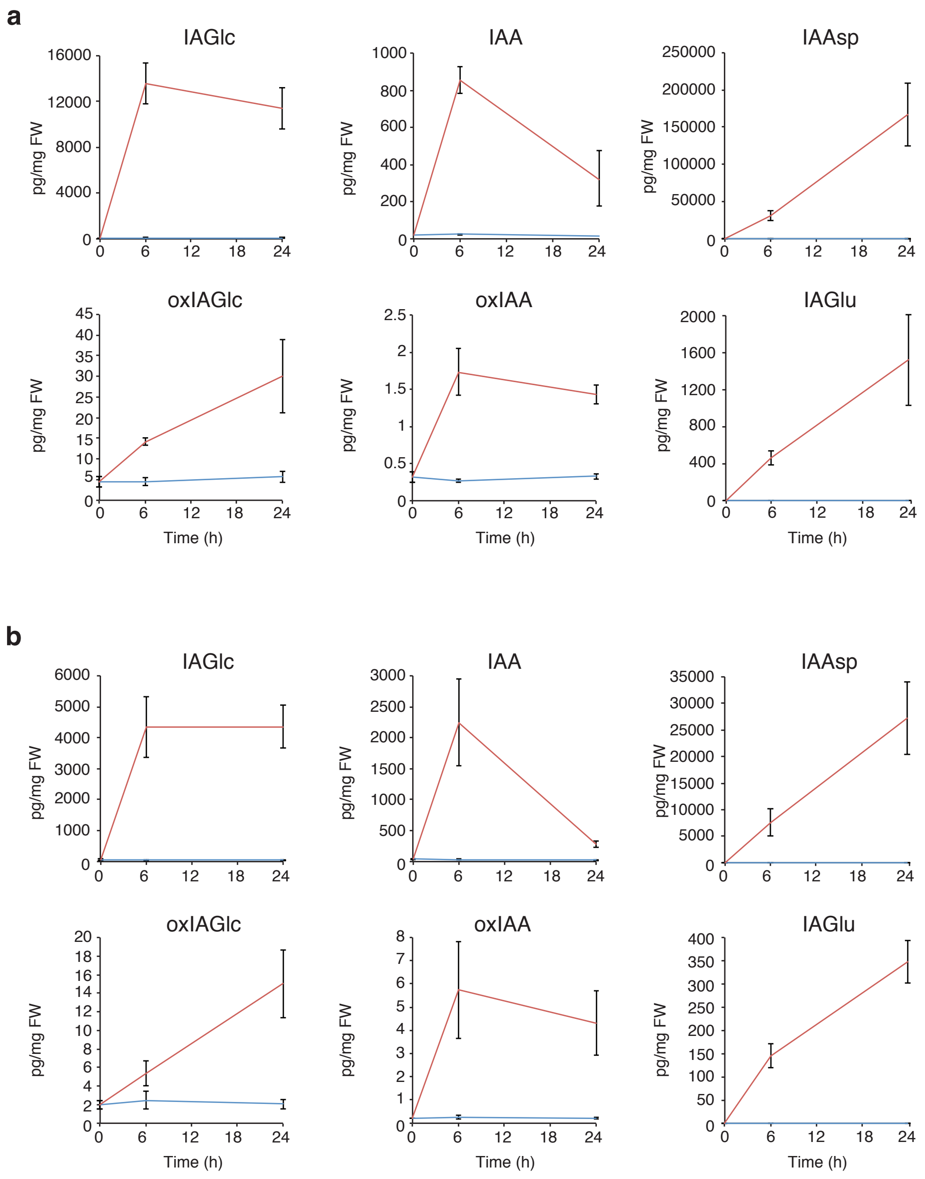


**Figure S4**

**Concentrations of IAA metabolites in *Pinus sylvetris* and *Pinus contorta* roots after feeding with unlabeled IAA.**

Two-week-old pine seedlings were incubated with (red line) or without (blue line) 10 $\mu$M unlabeled IAA and the levels of IAA and IAA metabolites oxIAA, oxIAGlc, IAGlc, IAAsp and IAGlu were quantified in roots after different incubation times. Panel **a** indicates *P. sylvestris* and panel **b** indicates *P. contorta*. The concentrations of all metabolites are in picograms per milligram fresh weight (FW). Error bars indicate ±SD (n = 4).

```
AtGH3.1         1 ------------------------------------------------------------
AtGH3.2         1 ------------------------------------------------------------
AtGH3.3         1 ------------------------------------------------------------
AtGH3.4         1 ------------------------------------------------------------
AtGH3.5         1 ------------------------------------------------------------
AtGH3.6         1 ------------------------------------------------------------
PpinGH3.16      1 ------------------------------------------------------------
PaGH3.16        1 ------------------------------------------------------------
PaGH3.gII.1     1 ------------------------------------------------------------
PaGH3.gII.2     1 ------------------------------------------------------------
PaGH3.gII.3     1 ------------------------------------------------------------
PaGH3.gII.4     1 ------------------------------------------------------------
PaGH3.gII.5     1 ------------------------------------------------------------
PaGH3.gII.6     1 ------------------------------------------------------------
PaGH3.gII.7     1 ------------------------------------------------------------
PaGH3.gII.8     1 ------------------------------------------------------------
PaGH3.gII.9     1 ------------------------------------------------------------
PaGH3.gII.10    1 ------------------------------------------------------------
AtGH3.9         1 ------------------------------------------------------------
AtGH3.17        1 ------------------------------------------------------------
PaGH3.17        1 ------------------------------------------------------------
AtGH3.11        1 ------------------------------------------------------------
AtGH3.10        1 ------------------------------------------------------------
PaGH3.gI.1      1 ------------------------------------------------------------
PaGH3.gI.2      1 MPGSIDSHEEVIREFEAMAQNAEEVIREFEAIAQNAERDNMKNRSEGGKRIEGEEGVKDE

AtGH3.1         1 ---------------------------------------------------------MAV
AtGH3.2         1 --------------------------------------------------------MAVD
AtGH3.3         1 ---------------------------------------------------------MTV
AtGH3.4         1 --------------------------------------------------------MAVD
AtGH3.5         1 -------------------------------------------------------MPEAP
AtGH3.6         1 -------------------------------------------------------MPEAP
PpinGH3.16      1 ---------------------------------------------------MPQAKREEG
PaGH3.16        1 ---------------------------------------------------MPQAKREES
PaGH3.gII.1     1 ------------------------------------------------------------
PaGH3.gII.2     1 ------------------------------------------------------------
PaGH3.gII.3     1 ------------------------------------------------------------
PaGH3.gII.4     1 ------------------------------------------------------------
PaGH3.gII.5     1 ------------------------------------------------------------
PaGH3.gII.6     1 ------------------------------------------------------------
PaGH3.gII.7     1 ------------------------------------------------------------
PaGH3.gII.8     1 ------------------------------------------------------------
PaGH3.gII.9     1 ------------------------------------------------------------
PaGH3.gII.10    1 ------------------------------------------------------------
AtGH3.9         1 ------------------------------------------------------------
AtGH3.17        1 ------------------------------------------------------------
PaGH3.17        1 ------------------------------------------------------------
AtGH3.11        1 -------------------------------------------------------MDALK
AtGH3.10        1 ------------------------------------------------------------
PaGH3.gI.1      1 ------------------------------------------------------------
PaGH3.gI.2     61 SRRSNMTGSVDSHEEVIRELEAMTLKAERVQRETLEDNMNKGSEGGKMIEGKEGALLNRV
```

```
AtGH3.1           4 DSNLSSPLGPPACEKDAKALRFIEEMTRNADTVQENLLAEILARNADTEYLRRFNLC---
AtGH3.2           5 SPLQSRMVSATTSEKDVKALKFIEEMTRNPDSVQEKVLGEILTRNSNTEYLKRFDLD---
AtGH3.3           4 DSALRSPMMHSPSTKDVKALRFIEEMTRNVDFVQKKVIREILSRNSDTEYLKRFGLK---
AtGH3.4           5 SLLQSGMASPTTSETEVKALKFIEEITRNPDSVQEKVLGEILSRNSNTEYLKRFDLN---
AtGH3.5           6 KKESLEVFDLTLDQKNKQKLQLIEELTSNADQVQRQVLEEILTRNADVEYLRRHDLN---
AtGH3.6           6 KIAALEVSDESLAEKNKNKLQFIEDVTTNADDVQRRVLEEILSRNADVEYLKRHGLE---
PpinGH3.16       10 IESVQESIIIATTERNRKALDFIEHATIHAAEVQAEVLLEILTRNAYTEYLERYQLT---
PaGH3.16         10 MESVQECISIATTERNKEALDFIENVTVHADAVQERVLLEILTRNAHTEYLQRYGFS---
PaGH3.gII.1       1 ----MSSVEKRASNREKKALEFIENITRNADEVQAQVLAAILTRNADTEYLKRYGLS---
PaGH3.gII.2       1 ------------------------------------------------------------
PaGH3.gII.3       1 ------------------------------------------------------------
PaGH3.gII.4       1 MPEAPMRMGNRANDRDKRALDFIENVTTNADDEQVRVLSAILTRNANTEYLKRYGLS---
PaGH3.gII.5       1 ------------------------------------------------------------
PaGH3.gII.6       1 ------------------------------------------------------------
PaGH3.gII.7       1 -------IGNGANDRHKAALEFIENVTRNADEVQARVLCSILTRNGDTEYLKRHGLS---
PaGH3.gII.8       1 ---MSSSLGNEVDDRNKRALEFIETVTTDADEVQTQVLSSILSRNADTEYLKRYGLN---
PaGH3.gII.9       1 ---MSSAMGNGVSDRDKRALDFIESVTTNADDVQIRVLSSILSRNADTEYLKRYGLN---
PaGH3.gII.10      1 ------------------------------------------------------------
AtGH3.9           1 --------MDVMKLDHDSVLKELERITSKAAEVQDNILRGILERNKDTEYLSKY-MN---
AtGH3.17          1 ------MIPSYDPNDTEAGLKLLEDLTTNAEAIQQQVLHQILSQNSGTQYLRAF-LD---
PaGH3.17          1 ----MEMNIRCAKDKGEAALQLIENLTARADEVQKQVLYEILRRNAETEYLNKF-LN---
AtGH3.11          6 HKRAFKMLEKVETFDMNRVIDEFDEMTRNAHQVQKQTLKEILKNQSAIYLQNCGLN---
AtGH3.10          1 --------METVEAGHDDVIGWFEHVSENACKVQSETLRRILELNSGVEYLRKWLGTVDV
PaGH3.gI.1        1 ------------------------------------------------------------
PaGH3.gI.2      121 KDESRRSDMPGSVDSHEEVIREFEAMTHNAERVQRETLEAILQRNSGTEYLKKWGVF---

AtGH3.1          61 GATDRDT----FKTKIPVITYEDLQPEIQRIADGDRSPILSAHPISEFLTSSGTSAGERK
AtGH3.2          62 GVVDR--KT--FKSKVPVVTYEDLKPEIQRISNGDCSPILSSHPITEFLTSSGTSAGERK
AtGH3.3          61 GFTDR--KT--FKTKVPVVIYDDLKPEIQRIANGDRSMILSSYPITEFLTSSGTSAGERK
AtGH3.4          62 GAVDR--KS--FKSKVPVVIYEDLKTDIQRISNGDRSPILSSHPITEFLTSSGTSAGERK
AtGH3.5          63 GRTDR--ET--FKNIMPVITYEDIEPEINRIANGDKSPILSSKPISEFLTSSGTSGGERK
AtGH3.6          63 GRTDR--ET--FKHIMPVVTYEDIQPEINRIANGDKSQVLCSNPISEFLTSSGTSGGERK
PpinGH3.16       67 GRTDR--KS--FKERLPVITYEDLQPEILRIANGDMSPILSAHPISEFLTSSGTSAGERK
PaGH3.16         67 GRTDR--ES--FKECFPVITYEDLQPEILRIANGDTSPILSAHPISEFLTSSGTSAGERK
PaGH3.gII.1      54 GQTDR--ET--FKKCLPVITYDDLKPEIHRIANGDTSPILSAHPISEFLLSTGTSAGQPK
PaGH3.gII.2       1 ------------------------------------------------------------
PaGH3.gII.3       1 ------------------------------------------------------------
PaGH3.gII.4      58 GRTDRST----FKKCLPVITYDDLKSEIHRIANGDTSPILSGHPVSEFLTSSGTSAGERK
PaGH3.gII.5       1 ------------------------------------------------------------
PaGH3.gII.6       1 ------------------------------------------------------------
PaGH3.gII.7      51 GRTNR--ST--FKKCLPVITYDDLKPEIRRIANGDTSPILSAHPVSEFLTSSGTSAGERK
PaGH3.gII.8      55 GRTDR--AT--FKKCLPVITYDDLKPEIRRIASGDTSPILSAHPVSEFLTSSGTSAGERK
PaGH3.gII.9      55 GRTDR--AT--FKECLPVITYDDLQPDIRRIVGRDTSPILSAHPVSEFLTSSGTSAGERK
PaGH3.gII.10      1 ------------------------------------------------------------
AtGH3.9          49 GSKDVLE----FKRAVPIIIYKDIYPYIQRIANGEDSSLITGHSITEILCSSGTSAGEPK
AtGH3.17         51 GEADKNQQS--FKNKVPVVNYDDVKPFIQRIADGESSDIVSAQPITELLTSSGTSAGKPK
PaGH3.17         53 GRIDN--KS--FKINVPVVNYEDIKPYIQRIANGDASAIISAEPISELLTSSGTSGGQPK
AtGH3.11         63 GNATDPEEA--FKSMVPLVTDVELEPYIKRMVDGDTSPILTGHPVPAISLSSGTSQGRPK
AtGH3.10         53 EKMDDYTLETLFTSLVPIVSHADLDPYIQRIADGETSPLLTQEPITVLSLSSGTTEGRQK
PaGH3.gI.1        1 ------------------------------------------------------------
PaGH3.gI.2      178 QGSTDPDPLQHFKSCVPVVCYPDLAPYMQRIADGDTSPILTADPITAFSLTAGTGDGNQK
```

```
AtGH3.1        117 LMPTIKEELD-----------------RRQLLYSLLMPVMNL-YVPGLDKGKGMYFLFVKS
AtGH3.2        118 LMPTIEEDLD-----------------RRQLLYSLLMPVMNL-YVPGLDKGKGLYFLFVKS
AtGH3.3        117 LMPTIDEDMD-----------------RRQLLYSLLMPVMNL-YVPGLDKGKALYFLFVKT
AtGH3.4        118 LMPTIEEDIN-----------------RRQLLGNLLMPVMNL-YVPGLDKGKGLYFLFVKS
AtGH3.5        119 LMPTIEEELD-----------------RRSLLYSLLMPVMSQ-FVPGLENGKGMYFLFIKS
AtGH3.6        119 LMPTIEEELD-----------------RRSLLYSLLMPVMDQ-FVPGLDKGKGMYFLFIKS
PpinGH3.16     123 IMPTIHEELK-----------------RRTLLYSLLMPVMNQ-YMKGLDKGKGMYFLFVKS
PaGH3.16       123 IMPTIHEELE-----------------RRTLLYSLLMPVMNQ-YMEGLDKGKGMYFLFIKS
PaGH3.gII.1    110 LVPTIEEDME-----------------RRALLISLVMPIMSQ-YMEGLDKGKGMYFLFINS
PaGH3.gII.2      1 -------------------------------------------------------------
PaGH3.gII.3      1 -------------------------------------------------------------
PaGH3.gII.4    114 LMPTIQEELE-----------------RKTLLYSLLMPVMNQ-------------------
PaGH3.gII.5      1 -------------------------------------------------------------
PaGH3.gII.6      1 -------------------------------------------------------------
PaGH3.gII.7    107 LMPTIQEEWE-----------------RRNLLYKLLMPVMNQ-YVQGLDKGKGMYFYFIKS
PaGH3.gII.8    111 LMPTIQEELE-----------------RKALLYSLLMPVMNQ-YMKGLDEGKGMYFFFIKS
PaGH3.gII.9    111 LMPTIQEELE-----------------RKALLYSLLMPVMNQ-YMQGLDKGKGMYFFFIKS
PaGH3.gII.10     1 -------------------------------------------------------------
AtGH3.9        105 LMPTIPEDLD-----------------RRTFLYNLIIPIVNK-YITGLDKGKAMYLNFVKA
AtGH3.17       109 LMPSTAEELE-----------------RKTFFYSMLVPIMNK-YVDGLDEGKGMYLLFIKP
PaGH3.17       109 IMPSIPEELIHRKTFLYNLLMPIMNKYRKTFLYNLLMPIMNK-YVPGLDKGKGMYLQFIKT
AtGH3.11       121 FIPFTDELME-----------------NTLQLFRTAFAFRNR-DFPIDDNGKALQFIFSSK
AtGH3.10       113 YVPFTRHSAQ-----------------TTLQIFRLSAAYRSR-FYPIREGGRILEFIYAGK
PaGH3.gI.1       1 -------------------------------------------LFPIKPGGRVLEFVYGSK
PaGH3.gI.2     238 LVPFNDSVIA-----------------SKLQYYRITNAYTARQFPPPQSQSRFLQFIYGSR

AtGH3.1        160 ETKTPGGLPARPVLTSYYKSEHFRSR--PYDPYNVYTSPNEAILCPDSFQSMYTQMLCGL
AtGH3.2        161 ESKTSGGLPARPVLTSYYKSDHFKRR--PYDPYNVYTSPNEAILCSDSSQSMYAQMLCGL
AtGH3.3        160 ESKTPGGLPARPVLTSYYKSEQFKRR--PNDPYNVYTSPNEAILCPDSSQSMYTQMLCGL
AtGH3.4        161 ESTTSGGLPARPALTSYYKSDYFRTS----DSDSVYTSPKEAILCCDSSQSMYTQMLCGL
AtGH3.5        162 ESKTPGGLPARPVLTSYYKSSHFKER--PYDPYTNYTSPNETILCSDSYQSMYSQMLCGL
AtGH3.6        162 ESKTPGGLPARPVLTSYYKSSHFKNR--PYDPYTNYTSPNQTILCSDSYQSMYSQMLCGL
PpinGH3.16     166 ETRTPGGLLARPVLTSYYKSQDFIER--PYDPYNVYTSPMEAILCSDSYQSMYCQLLCGL
PaGH3.16       166 ETRTPGGLLARPVLTSYYKSQHFRER--PYDPYNVYTSPIEAILSADSYQSMYCQLLCGL
PaGH3.gII.1    153 ESKTPGGLLARYSSTRWYKSRFLKDKPLPYDPYNVYTSPIETILCPDAYQSMYCQLLCGL
PaGH3.gII.2      1 ------------------------------------------------------------
PaGH3.gII.3      1 ------------------------------------------------------------
PaGH3.gII.4        ------------------------------------------------------------
PaGH3.gII.5      1 ----------------------------------------------------MCD-----
PaGH3.gII.6      1 ------------------------------------------------------------
PaGH3.gII.7    150 EAKTPCGLLARPVLTSYYKSHYFRER--PYDPYNVCTSPIQTILCPDAYQSMYSQLLCGL
PaGH3.gII.8    154 EAKTPGGLLARPVLTSYYKSDYFKER--PYDPYNVYTSPNQTVLCQDAYQSMYSQLLCGL
PaGH3.gII.9    154 EAKTPGGLPARPVLTSYYKSPHFRER--PYDPYNVYTSPNQTVLCPDAYQSMYSQLLCGL
PaGH3.gII.10     1 ------------------------------------------------------------
AtGH3.9        148 ETSTPCGLPIRAVLTSYYKSKHFQCR--PYDPFNDLTSPIQTILCEDSNQSMYCQLLAGL
AtGH3.17       152 EIKTPSGLMARPVLTSYYKSQHFRNR--PFNKYNVYTSPDQTILCQDSKQSMYCQLLCGL
PaGH3.17       168 EVTTPSGLKARPVLTSYYKSSNFRDR--PFDKFNVYTSPDETILCPDSRQSMFCQLLCGL
AtGH3.11       164 QYISTGGVPVGTATTNVYRNPNFKAG--MKSITSPSCSPDEVIFSPDVHQALYCHLLSGI
AtGH3.10       156 EFKTLGGLTVGTATTHYYASEEFKTK--QETTKSFTCSPQEVISGGDFGQCTYCHLLLGL
PaGH3.gI.1      19 ESSTKGGLVASTATTNIYRSESFKIY--KKNIQILGCSPDEVIFGFDSRQSMYCHILCGL
PaGH3.gI.2     282 QFLTKGGLEASNASGLGLRSKGFKKY--KESNQWLACSPEEVVFGFNYEQSLYCHLLCGL
```

```
AtGH3.1        218 LDRLSVLRVGAVFASGLLRAIRFLQLHWSRFAHDIELGCLDS-EITDPSIRQCMSG-ILK
AtGH3.2        219 LMRHEVLRLGAVFASGLLRAISFLQNNWKELARDISTGTLSS-RIFDPAIKNRMSKILTK
AtGH3.3        218 LMRHEVLRLGAVFASGLLRAIGFLQTNWKELADDISTGTLSS-RISDPAIKESMSKILTK
AtGH3.4        217 LMRHEVNRLGAVFPSGLLRAISFLQNNWKELSQDISTGTLSS-KIFDHAIKTRMSNILNK
AtGH3.5        220 CQHQEVLRVGAVFASGFIRAIKFLEKHWIELVRDIRTGTLSS-LITDPSVREAVAK-ILK
AtGH3.6        220 CQHKEVLRVGAVFASGFIRAIKFLEKHWPELARDIRTGTLSS-EITDSSVREAVGE-ILK
PpinGH3.16     224 AQNHEVLRVGAVFASGLLRAIRFLEEHWQSLCQDIRSGTVNDEEVTDPCLRESVMK-ILH
PaGH3.16       224 AQNHEVLRVGAVFASGLLRAIRFLEEHWKSLCQDIRSGTINDEEVTDPCLRESVMK-ILR
PaGH3.gII.1    213 LQNYEVLRMGAIFASGFIRTIRFLEEHWRQLCLDMKTGILNK-EVTDPSVRETVEKLLLH
PaGH3.gII.2      1 ------------------------------------------------------------
PaGH3.gII.3      1 ------------------------------------------------------------
PaGH3.gII.4        ------------------------------------------------------------
PaGH3.gII.5      4 ------------------------------------------------------------
PaGH3.gII.6      1 ------------------------------------------------------------
PaGH3.gII.7    208 LQNNE-------------------------------------------------------
PaGH3.gII.8    212 LQNNEVLRMGAVFASGFIRAIRFLEEHWRQFCLDIKTGILNR-EVTDPSVREAVGE-LLH
PaGH3.gII.9    212 LQHNEVLRMGAVFASGFIRAIRFLEEHWTQFCLDIRTGIINR-EVTDPLVREAVGE-LLH
PaGH3.gII.10     1 ------------------------------------------------------------
AtGH3.9        206 IHRHKVMRLGAVFASAFLRAISYLEKKWSQLCEDIRTGSLNP-MITDPGCQMAMSCLLMS
AtGH3.17       210 VQRSHVLRVGAVFASAFLRAVKFLEDHYKELCADIRTGTVTS-WITDSSCRDSVLSILNG
PaGH3.17       226 LQRDEVLRVGAVFASAFLRAIKFLEENWEELCDNIRTGHLSD-WIDDPPSRIAVMK-MLS
AtGH3.11       222 LFRDQVQYVFAVFAHGLVHAFRTFEQVWEEIVTDIKDGVLSN-RITVPSVRTAMSK-LLT
AtGH3.10       214 HYSSQVEFVASAFSYTIVQAFSFFEEIWREICADIKEGNLSS-RITLPKMRKAVLA-LIR
PaGH3.gI.1      77 LYSNEVQFMSSTFSYSIVEAFRTFEQDWQQLCNDIKEGKLNE-KITVPSMRASVSK-LLK
PaGH3.gI.2     340 LYSYEVERLTSSFAYSIVEAFRTFEGVWQQLCTDIKEGTLNK-EITVPSMRESVSK-ILK

AtGH3.1        276 PDPVLAEFIRRECKSD--N--WEKIITRIWPNTKYLDVIVTGAMAQYIPTLEYYS-GGLP
AtGH3.2        278 PDQELAEFLVGVCSQE--N--WEGIITKIWPNTKYLDVIVTGAMAQYIPTLEYYS-GGLP
AtGH3.3        277 PDQELADFITSVCGQD--N-SWEGIITKIWPNTKYLDVIVTGAMAQYIPMLEYYS-GGLP
AtGH3.4        276 PDQELAEFLIGVCSQE--N--WEGIITKIWPNTKYLDVIVTGAMAEYIPMLEYYS-GGLP
AtGH3.5        278 PSPKLADFVEFECKKS--S--WQGIITRLWPNTKYVDVIVTGTMSQYIPTLDYYS-NGLP
AtGH3.6        278 PDPKLADFVESECRKT--S--WQGIITRLWPNTKYVDVIVTGTMSQYIPTLDYYS-NGLP
PpinGH3.16     283 PNTQLADLIRTECSKE--S--WQGIITRLWPNARYLDVIVTGAMAQYIKTLDFYS-GGLP
PaGH3.16       283 PKIQLADLIHAECSKE--S--WQGIITRLWPNARYLDVIVTGAMAQYIETLDFYS-GGLP
PaGH3.gII.1    272 PNPELAELIQRNCSAP--S--WQGIITRLWPNTKYIKTVVTGAMAQYIPTLDYYS-GGLP
PaGH3.gII.2      1 ------------------------------------------------------------
PaGH3.gII.3      1 ------------------------------------------------------------
PaGH3.gII.4        ------------------------------------------------------------
PaGH3.gII.5      4 ---------------------W------------------------------------FP
PaGH3.gII.6      1 ------------------------------------------------------------
PaGH3.gII.7    213 ----------------------------------------------YIPTLDYYS-AGLP
PaGH3.gII.8    270 PNPELADFVERKCSAQ--S--WQGIITRLWPNTKYIDVIVTGAMAQYIPTLDYYS-GGLP
PaGH3.gII.9    270 PNPELADLVERECSAP--S--WQGIIARLWSNTKYIDVIVTGAMAQYIPTLDYYS-GGLP
PaGH3.gII.10     1 ------------------------------------------------------------
AtGH3.9        265 PNPELASEIEEICGRS--S--WKGILCQLWPKAKFIEAVVTGSMAQYIPALEFFSQGKIP
AtGH3.17       269 PNQELADEIESECAEK--S--WEGILRRIWPKAKYVEVIVTGSMAQYIPTLEFYS-GGLP
PaGH3.17       284 PNPQLAEEIHGECSKK--S--WQGIITRLWRKTIYIDVIVTGTMAQYIPTLDYYG-GGLP
AtGH3.11       280 PNPELAETIRTKCMSL--S-NWYGLIPALFPNAKYVYGIMTGSMEPYVPKLRHYA-GDLP
AtGH3.10       272 PNPSLASHIEEICLELETNLGWFGLISKLWPNAKFISSIMTGSMLPYLNKLRHYA-GGLP
PaGH3.gI.1     135 PDPDLADAIYNKCKNI--G-NWHGVIPLLWPNAKYIISIMTGAMEPYLRKLRHYA-GDLP
PaGH3.gI.2     398 PNPELAQSIFETCEKLLTN-NWYGVIHKLWPNVKYIGCIITGSMEAYVKKLRHYA-GSLP
```

```
AtGH3.1       331 MACTMYASSECYFGLNLNPMSKPSEVSYTIMPNMAYFEFIPLG-----------------
AtGH3.2       333 MACTMYASSESYFGINLKPMCKPSEVSYTIMPNMAYFEFLPHNHDGD-----------GA
AtGH3.3       333 MACTMYASSESYFGINLKPMCKPSEVSYTIMPNMAYFEFLPHHEVPTE------------
AtGH3.4       331 MASMIYASSESYFGINLNPMCKPSEVSYTIFPNMAYFEFLPHNHDGDG------------
AtGH3.5       333 LVCTMYASSECYFGVNLRPLCKPSEVSYTLIPSMAYFEFLPVHRNNG----VTNSINLPK
AtGH3.6       333 LVCTMYASSECYFGVNLRPLCKPSEVSYTLIPNMAYFEFLPVHRNSG----VTSSISLPK
PpinGH3.16    338 QVCTMYASSECYFGINLKPLCQPWEVSYTLMPNMAFFEFLPVYRNKNEDAG-------PV
PaGH3.16      338 QVCTMYASSECYFGINLKPLCKSSEVSYTLMPNMAFFEFLPVYRNNDDA---------AP
PaGH3.gII.1   327 LVCTMYSSSEGSFGLNLNPLCKPTEVSYTLLPNMAYFEFLPVHRQEFG--LSIESPAIPK
PaGH3.gII.2     1 ---------------------------------MAYFEFLPVHRKQEEAGATLESPAITK
PaGH3.gII.3     1 ------------------------------------------------------------
PaGH3.gII.4       ------------------------------------------------------------
PaGH3.gII.5     7 QRCA--------------------------------------------------------
PaGH3.gII.6     1 ------------------------------------------------------------
PaGH3.gII.7   226 LVCTMYASSECYFGLNLKPLCKPSEVSYTLLPNMAYFEFLPVHRKLQLEAQE------CP
PaGH3.gII.8   325 LVCTMYASSECYFGLNLKPLCKPSEVSYTLLPNMAYFEFLPVHRKQEAAGLTIESSTIPK
PaGH3.gII.9   325 LVCTMYASSECYFGLNLKPLCKPFEVSYTLLPNMAYFEFLPVHRKQEEAGVTLES---PG
PaGH3.gII.10    1 ------------------------------------------------------------
AtGH3.9       321 LVCPMYASSETYFGVNVEPLSKPSDVVFTLLPNMCYFEFIPLGKNGTLSFD-------LD
AtGH3.17      324 LVSTMYASSECYFGINLNPLCDPADVSYTLLPNMAYFEFLPVDDKSHEEIHFATHSNTDD
PaGH3.17      339 LVSTMYASSECYFGVNLKPL-----------------KVLDLQSNGTE------------
AtGH3.11      336 LVSHDYGSSEGWIAANVTPRLSPEEATFAVIPNLGYFEFLPVSETGEG------------
AtGH3.10      331 LVSADYGSTESWIGVNVDPHLPPEDVSFAVIPTFSYFEFIPLYRRQNQSDICI-------
PaGH3.gI.1    191 LLTSEYGATEGWIAANIDPTSSPENTIYTVVPDIGYFEFIPLHLRHEGLKLDN-----SV
PaGH3.gI.2    456 ILPGGYAVSEGYLAVNIDS----ATTTFTVVPSTAFFEFIPQDKEG--------------

AtGH3.1       374 ------GTKAVELVDVNIGKEYELVVTTYAGLCRYRVGDILRVTGFHNSAPQFHFVRRKN
AtGH3.2       382 AEASLDETSLVELANVEVGKEYELVITTYAGLYRYRVGDILRVTGFHNSAPQFKFIRRKN
AtGH3.3       381 ------KSELVELADVEVGKEYELVITTYAGLNRYRVGDILQVTGFYNSAPQFKFVRRKN
AtGH3.4       379 ---GVEATSLVELADVEVGKEYELVITTYAGLYRYRVGDILRVTGFHNSAPQFKFIRREN
AtGH3.5       389 ALTEKEQQELVDLVDVKLGQEYELVVTTYAGLCRYRVGDLLRVTGFKNKAPQFSFICRKN
AtGH3.6       389 ALTEKEQQELVDLVDVKLGQEYELVVTTYAGLYRYRVGDVLSVAGFKNNAPQFSFICRKN
PpinGH3.16    391 TTATEQPAELVDLVDVKVGQEYELVITTYSGLYRYRVGDVLRVTGFHNAAPQFQFVCRKN
PaGH3.16      389 VTMATEQQELVDLADVTVGQEYELVITTYAGLYRYRVGDVLRVTGFYNAAPQFQFVCRKN
PaGH3.gII.1   385 AVDEKEEKDLVKLVDVKVGQEYELVVTTYAGLYRYRVGDVLRVTGFHNAAPQFHFVCRQN
PaGH3.gII.2    28 TLDEKERQELVELVDVKLGQEYELVVTTYAGLNRYRVGDVLRVTGFYNAAPQFHFVCRQN
PaGH3.gII.3     1 ------------------------------------------------------------
PaGH3.gII.4       ------------------------------------------------------------
PaGH3.gII.5    11 -----------------------------------------------------SIPCRQN
PaGH3.gII.6     1 ------------------------------------------MTGFHNTAPQFHFVYRQN
PaGH3.gII.7   280 TLNEKDQDELLDLVDVKIGQEYELVITTYAGLYRYRVGDVLRVTGFHNAAPQFHFVCRQN
PaGH3.gII.8   385 TLDDKEREELVELVDLKLGQEYELVVTTYAGLNRYRVGDVLRVTGFHNAAPQFHFVCRQN
PaGH3.gII.9   382 TLDEKEREELVELADVKLGREYELVVTTYAGLNRYRVGDVLRVTGFYNAAPQFHFVCRQN
PaGH3.gII.10    1 ---------------------------------------------MHNS----NFVCRQN
AtGH3.9       374 DDEQVPCDKVVDLVNVKLGRYYELVVTTFAGLYRYRIGDVLQVAGFYNGAPQFRFICRRN
AtGH3.17      384 DDDALKEDLIVNLVNVEVGQYYEIVITTFTGLYRYRVGDILKVTGFHNKAPQFRFVQRRN
PaGH3.17      370 -NGKKIEDELVDLVDVKVGHYYELVVTTYAGLYRYRVGDILLVTGFYNRAPQFEFVYRRN
AtGH3.11      384 ------EEKPVGLTQVKIGEEYEVVITNYAGLYRYRLGDVVKVIGFYNNTPQLKFICRRN
AtGH3.10      384 -DGDFVEDKPVPLSQVKLGQEYELVLTTFTGLYRYRLGDVVEVTSFHKGTPKLSFIYRRK
PaGH3.gI.1    246 ATADYIESEPVGLTEVKVGQEYEVVLTTFAGLYRYRLGDVVKVTGFYNSSPQLSYICRKN
PaGH3.gI.2    498 ------QAEPIGLTEVEIGKEYEVVITTFGG-----------------------------
```

```
AtGH3.1       428 VLLSIDSDKTDESELQKAVENAS--SILHEECGSRVAEYTSYADTSTIPGHYVLYWELLV
AtGH3.2       442 VLLSVESDKTDEAELQKAVENAS--RLFAEQ-GTRVIEYTSYAETKTIPGHYVIYWELLG
AtGH3.3       435 VLLSIESDKTDEAELQSAVENAS--LLLGEQ-GTRVIEYTSYAETKTIPGHYVIYWELLV
AtGH3.4       436 VLLSIESDKTDEADLQKAVENAS--RLLAEQ-GTRVIEYTSYADTKTIPGHYVIYWELLS
AtGH3.5       449 VVLSIDSDKTDEVELQNAVKNAV--THLVPF-DASLSEYTSYADTSSIPGHYVLFWELCL
AtGH3.6       449 VVLSIDSDKTDEVELQNAVKNAV--THLVPF-DASLSEYTSYADTSSIPGHYVLFWELCL
PpinGH3.16    451 VMLSIDADKTDEAELHNAVMNAV--KHLEPL-EATLVEYTSYTDTSTIPGHYVLYWELRT
PaGH3.16      449 VMLSIDSDKTDEAELHSAVKNAV--KHLEPF-EASLVEHTSYAETSTIPGHYVLYWELRN
PaGH3.gII.1   445 VVLSIHIDKTDEAELQSSVEKSI--KHLKPF-DITLMDYTSYADTSTIPGHYVLYWELRF
PaGH3.gII.2    88 VVLSIDADKTDEAELHSAVENSV--KHLEPF-DAQLIEYTSYADTSTIPGHYVLYWELRF
PaGH3.gII.3     1 ---------MDEAELHSAVENSA--KHLEPF-DAQLIEYTSYADTATIPGHYMLYWELHF
PaGH3.gII.4       ------------------------------------------------------------
PaGH3.gII.5    18 VVLSIDSDKTDEAERHSAVENSA--KHLEPF-DAQLIEYTRYSDTATIPGHYVLYWELHF
PaGH3.gII.6    19 VVISIDSDKRDEVELHSAVENFA--KHLEPF-NAQLIEYTSYADIATIPGHYVLYWELRF
PaGH3.gII.7   340 VVLSIDSDKTDEAELQSAVENSV--KHLEPF-DSKLIEYTSYADTSTIPGHYVLYWELGS
PaGH3.gII.8   445 VVLSIDSDKTDEVELHSAVENSV--KHLEAF-DAQLIEYTSYADTGTIPGHYVLYWELRF
PaGH3.gII.9   442 VVLSIDSDKTDEAELHSAVENSA--KHLVPF-HAQLVEYTSYADTATIPGHYVLYWELRF
PaGH3.gII.10   12 VVLSIDSNKTDEVELHSAIENSV--KHLEAF-DAQLIEYTSYAETGTIPGHYVLYWELRF
AtGH3.9       434 VVLSIDLDKTNEEDLHRSITLAK--KKLGS--NAFLAEYTSYADTSSVPGHYVLFWEIQG
AtGH3.17      444 VVLSIDTDKTSEEDLLNAVTQAKLNHLQHPS-SLLLTEYTSYADTSSIPGHYVLFWELKP
PaGH3.17      429 VVLSIDTDKTNEEDLLKAVTKAK--KLLEPF-NALLSEYTSYADTSTLPGHYVLFWELNT
AtGH3.11      438 LILSINIDKNTERDLQLSVESAA--KRLSEE-KIEVIDFSSYIDVSTDPGHYAIFWEISG
AtGH3.10      443 LILTINIDKNTEKDLQRVVDKAS--QLLSRSTRAEVVDFTSHADVIARPGHYVIYWEIRG
PaGH3.gI.1    306 LILTVNIDKTTEKDLQISVDKAT--ELLKEE-NVDLVDFTSYADLSTVPGHYVIFWELSD
PaGH3.gI.2    523 ----------------------------------------------------AFYINL--

AtGH3.1       486 RDGA---RQPSHETLTRCCLGMEESLNSVYRQSRVADNSVGPLEIRVVRNGTFEELMDYA
AtGH3.2       499 RDQSN--ALMSEEVMAKCCLEMEESLNSVYRQSRVADKSIGPLEIRVVRNGTFEELMDYA
AtGH3.3       492 KDQT---NPPNDEVMARCCLEMEESLNSVYRQSRVADKSIGPLEIRVVKNGTFEELMDYA
AtGH3.4       493 RDQSN--ALPSDEVMAKCCLEMEESLNAVYRQSRVSDKSIGPLEIRVVQNGTFEELMDFS
AtGH3.5       506 DGNT---PIPP-SVFEDCCLAVEESFNTVYRQGRVSDKSIGPLEIKIVEPGTFDKLMDYA
AtGH3.6       506 NGNT---PIPP-SVFEDCCLTIEESLNSVYRQGRVSDKSIGPLEIKMVESGTFDKLMDYA
PpinGH3.16    508 SAL----PVPP-SVFEDCCLTVEESLNSVYRQCRVADKSIGPLEIKVVEMGTFDKLMDYA
PaGH3.16      506 STV----PVPA-SVFEDCCLTIEESLNSVYRQCRVADKSIGPLEIKVVETGTFDKLMDYA
PaGH3.gII.1   502 NNKEVAGAVPS-SVFEDCCLTVEESLNFVYRQGRGADKSIAPLEIRVVEEGTFEQLMDRA
PaGH3.gII.2   145 NTKSVADAVPS-SVFEDCCLTVEESLNSVYRQGRASDNSIA-------------------
PaGH3.gII.3    49 NTKTV--EVPF-SVFEDCCLTAEESLNSIYRQGRASDKSIGPMEIRVVEEGMFDELMDYA
PaGH3.gII.4       ------------------------------------------------------------
PaGH3.gII.5    75 STKTV--EVPF-SVFEDCCLTAEESLNSIYRQGRASDKSIDPMEIRVVEEETFDELMDYA
PaGH3.gII.6    76 NTKAV--EVPS-LVFEDCCLTTEESLNFVYRQGRAFDKSIGPLEIRVVEEGTFDELMDYA
PaGH3.gII.7   397 KEVAL--TIPP-SVFEDCCLTVEESLNSVYRQGRVSDKSIGPLEIRVVEDGTFDELMDYA
PaGH3.gII.8   502 NTKAI--EVPS-SVFEDCCLTAEESLNSVYRQGRASDKSIGPLEIRVVEEGTFNELMDYA
PaGH3.gII.9   499 DTKAV--AFPS-SVFEDCCLTIEESLNSVYRQGRASDRSIGPLEIRVVEEGTFDQLMDYA
PaGH3.gII.10   69 NTKAI--EVPS-SVFEDCCLTAEESLNSVYRQGRASDKSIGPLEIRVVEEGTFDELMDYA
AtGH3.9       490 HLEP--------KLMEECCVAVEEELDYIYRQCRTKERSIGALEIRVVKPGTFEKLMDLI
AtGH3.17      503 RHSNDPPKLDD-KTMEDCCSEVEDCLDYVYRRCRNRDKSIGPLEIRVVSLGTFDSLMDFC
PaGH3.17      486 RE-----EFLDASVLESCCSTIEESLDSIYRRCRTKDKSIGPLEIRLVKPGTFDLLMDYC
AtGH3.11      495 ETNE--------DVLQDCCNCLDRAFIDAGYVSSRKCKTIGALELRVVAKGTFRKIQEHF
AtGH3.10      501 EADD--------KALEECCREMDTAFVDYGYVVSRRMNSIGPLELRVVERGTFGKVAERC
PaGH3.gI.1    363 SLNE--------GIVKRCCSIMDETFIDPGYVVSRKANTIGPLELRIVERGTFRKIIDYY
PaGH3.gI.2        ------------------------------------------------------------
```

```
AtGH3.1       543 ISRGASINQYKVPRCVN-F-TPIVELLDSRVVSAHFSPSLPHWTPERRRR--------
AtGH3.2       557 ISRGASINQYKVPRCVS-F-TPIMELLDSRVVSAHFSPSLPHWSPERRR---------
AtGH3.3       549 ISRGASINQYKVPRCVS-F-TPIMELLDSRVVSTHFSPALPHWSPERRR---------
AtGH3.4       551 ISRGSSINQYKVPRCVS-L-TPIMKLLDSRVVSAHFSPSLPHWSPERRH---------
AtGH3.5       562 ISLGASINQYKTPRCVK-F-APIIELLNSRVVDSYFSPKCPKWVPGHKQWGSN-----
AtGH3.6       562 ISLGASINQYKTPRCVK-F-APIIELLNSRVVDSYFSPKCPKWSPGHKQWGSN-----
PpinGH3.16    563 ISRGSSINQYKAARCVK-F-APMVDILNSRVSASYFSPRCPKWTAGRTQWGALPR---
PaGH3.16      561 ISRGSSINQYKAPRCVK-F-APMVDLLKSRVSASYFSPRCPKWTPGRTQWGAITRLI-
PaGH3.gII.1   561 VCQKASMNQYKAPRCVK-S-TSMVELLNSRVIHSYFSPRSPMWVPGSAVSKLVGS---
PaGH3.gII.2       ----------------------------------------------------------
PaGH3.gII.3   106 LSRGASINQYKAPRCIK-F-APIVELLNSRVVHSYFNPHNGLSDAEIGHNISLLK---
PaGH3.gII.4       ----------------------------------------------------------
PaGH3.gII.5   132 LSRGASINQ-------------------------------------------------
PaGH3.gII.6   133 E---------------------------------------------------------
PaGH3.gII.7   454 LSRGASINQYKVPRCVK-C-SPIVGLLNSRVMHSYFSPKCPQWAPGCRKWA-------
PaGH3.gII.8   559 LSRGASINQYKAPRCIK-F-TPIVELLNSRVVHSYFSPQPPQWAPGCRNWA-------
PaGH3.gII.9   556 LSRGASINQYKAPRCVK-F-TPIVELLNSRVVHSYFSTQPPQWAPGCREWA-------
PaGH3.gII.10  126 LSRGASINQYKAPRCIK-F-TPIVELLNSRVVHSYFSPQPPQWAPGCRNWA-------
AtGH3.9       542 ISQGGSFNQYKTPRCVK-SNSATFKLLNGHVMASFFSPRDPTWVP--------------
AtGH3.17      562 VSQGSSLNQYKTPRCVK-S-GGALEILDSRVIGRFFSKRVPQWEPLGLDS--------
PaGH3.17      541 LNQGSSYNQYKTPRCIK-S-LHVLGLLNSKVTAKYFSKRLPSWTPYNPGSLNPNSQLY
AtGH3.11      547 LGLGSSAGQFKMPRCVKPSNAKVLQILCENVVSSYFSTAF------------------
AtGH3.10      553 VGKCGGLNQFKTPRCT--TNSVMLDILNDSTIKRFRSSAYD-----------------
PaGH3.gI.1    415 TSHGGAVNQFKTPRCIS-TNQALLDILNKNTIQTCFSTLFS-----------------
PaGH3.gI.2        ----------------------------------------------------------
```

**Fig. S5**

**Multiple sequence alignment of predicted amino acid sequences of PaGH3 and AtGH3 proteins.**

The multiple sequence alignment was obtained with MUSCLE software. Black shading indicates conserved residues, gray shading indicates residues with similar properties, - indicates a gap inserted to maximize the alignment. Letters with different colors indicate motif 1 (red), motif 2 (blue) and motif 3 (orange) involved in ATP/AMP binding that are conserved in the acyl-adenylate/thioesther-forming enzyme superfamily (Staswick *et al.*, 2002). Bold letters represent conserved motif variation in members of Group I.

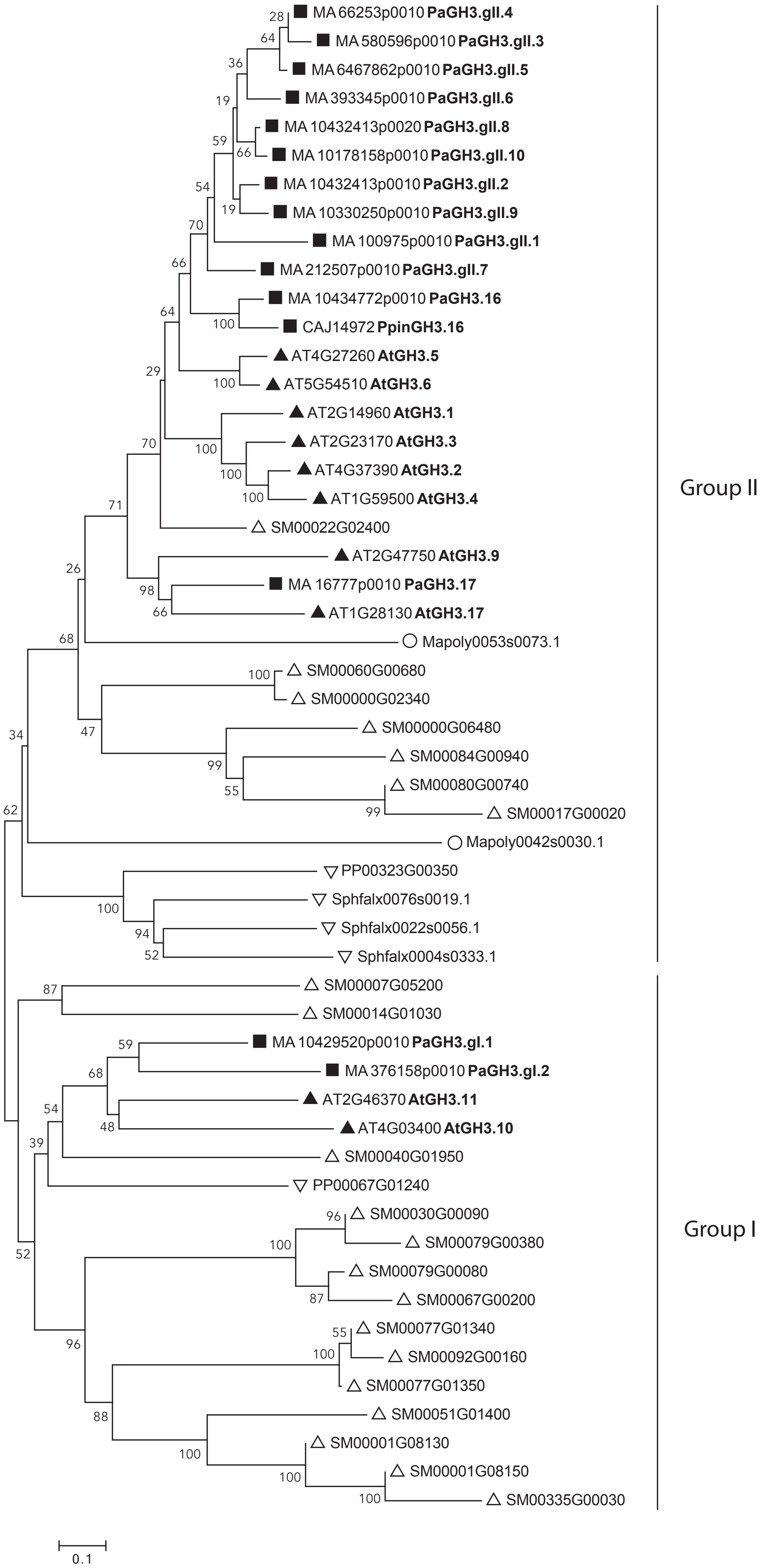
MA 66253p0010 PaGH3.gII.4
MA 580596p0010 PaGH3.gII.3
MA 6467862p0010 PaGH3.gII.5
MA 393345p0010 PaGH3.gII.6
MA 10432413p0020 PaGH3.gII.8
MA 10178158p0010 PaGH3.gII.10
MA 10432413p0010 PaGH3.gII.2
MA 10330250p0010 PaGH3.gII.9
MA 100975p0010 PaGH3.gII.1
MA 212507p0010 PaGH3.gII.7
MA 10434772p0010 PaGH3.16
CAJ14972 PpinGH3.16
AT4G27260 AtGH3.5
AT5G54510 AtGH3.6
AT2G14960 AtGH3.1
AT2G23170 AtGH3.3
AT4G37390 AtGH3.2
AT1G59500 AtGH3.4
SM00022G02400
AT2G47750 AtGH3.9
MA 16777p0010 PaGH3.17
AT1G28130 AtGH3.17
Mapoly0053s0073.1
SM00060G00680
SM00000G02340
SM00000G06480
SM00084G00940
SM00080G00740
SM00017G00020
Mapoly0042s0030.1
PP00323G00350
Sphfalx0076s0019.1
Sphfalx0022s0056.1
Sphfalx0004s0333.1
SM00007G05200
SM00014G01030
MA 10429520p0010 PaGH3.gI.1
MA 376158p0010 PaGH3.gI.2
AT2G46370 AtGH3.11
AT4G03400 AtGH3.10
SM00040G01950
PP00067G01240
SM00030G00090
SM00079G00380
SM00079G00080
SM00067G00200
SM00077G01340
SM00092G00160
SM00077G01350
SM00051G01400
SM00001G08130
SM00001G08150
SM00335G00030
Group II
Group I
0.1

**Fig. S6**

**Phylogenetic relationships of GH3 proteins between *Picea abies* and other land plants.**

Predicted proteins sequences (both full-length and partial) from *P. abies* (Pa), *Pinus pinaster* (Ppin), Arabidopsis (AT), *Selaginella moellendorffii* (SM), *Physcomitrella patens* (PP), *Sphagnum fallax* (Sphfalx) and *Marchantia polymorpha* (Mapoly) were aligned using MUSCLE program. The phylogenetic tree was constructed using MEGA6 program and the Neighbor-Joining method with predicted GH3 proteins. Bootstrap support is indicated at each node. Closed triangle indicates angiosperms, closed square represents gymnosperms, up-pointing open triangle indicates lycophytes, down-pointing open triangle represents mosses and open circle indicates liverworts.

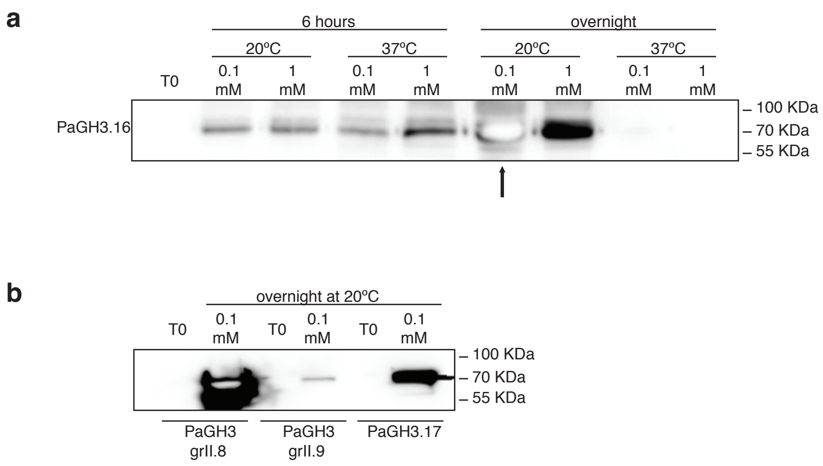


**Figure S7**

**Western blot analysis of IPTG-induced PaGH3.16, PaGH3.gII.8, PaGH3.gII.9 and PaGH3.17 recombinant proteins.**

**(a)** PaGH3.16 was used to test the optimal condition for expression of PaGH3 proteins. Bacterial cultures were treated with either 0.1 or 1 mM IPTG and incubated at 20°C or 37°C for 6 hours or overnight. Bacterial cultures that were sampled before the IPTG induction were used as non-induced control (at time 0, T0). The optimal condition was set to 0.1 mM IPTG overnight at 20°C (indicated by a black arrow) and used to induce the expression of other PaGH3 proteins. **(b)** PaGH3.gII.8, PaGH3.gII.9 and PaGH3.17 proteins expression. Calculated molecular weight (MW) of 6x His-recombinant proteins: 70.84 kDa (PaGH3.16),
